# Inflammatory memory disables a DDR1 degradation checkpoint to enable pancreatic cancer growth

**DOI:** 10.64898/2026.04.01.715820

**Authors:** Fei Yang, Bo Lin, Zihang Yuan, Yongkang Yuan, Xiaohong Pu, Chunlan Wang, Kosuke Watari, Rongkui Luo, Beicheng Sun, Michael Karin, Hua Su

## Abstract

Inflammation and stroma remodeling regulate pancreatic ductal adenocarcinoma (PDAC), but how or if these cues are integrated at the molecular level remains unclear. Here, we identify a metabolic checkpoint that controls the stability of the collagen receptor DDR1, and subsequent tumorigenesis. We show that defective Col-I remodeling deprives PDAC cells of the high affinity DDR1 ligand, ¾Col-I, resulting in reduced ATP and activation of AMPK. AMPK phosphorylates DDR1 at T519, promoting its recognition by the E3 ubiquitin ligase adaptor FBXW2 and subsequent degradation. Importantly, this degradation pathway can be disabled by inflammatory signaling. Exposure to inflammatory cytokines induces methylation-dependent silencing of FBXW2, which establishes an inflammatory memory that preserves DDR1 stability, enabling sustained ligand-triggered receptor oligomerization and downstream NF-κB-NRF2 signaling even in restrictive stromal environments. Together, these findings identify regulated receptor turnover as a mechanism through which stromal architecture, metabolic state, and inflammatory memory are integrated to control PDAC progression.

## Introduction

Pancreatic ductal adenocarcinoma (PDAC) is one of the deadliest human malignancies defined by a dense desmoplastic stroma dominated by type I collagen (Col-I) fibrils and chronic inflammation that profoundly shape tumor cell behavior (1–3). Once considered a mere structural scaffold, Col-I has emerged as a dynamic signaling and metabolic regulator of the PDAC tumor microenvironment (TME), imposing selective pressures that require adaptive programs for tumor growth and survival (4–6). However, whether and how PDAC cells integrate stromal architecture with inflammatory cues to establish such adaptive states remains unclear.

Proteolytic remodeling by matrix metalloproteases (MMPs) generates soluble ¾ collagen fragments that promote PDAC progression by activating discoidin domain receptor 1 (DDR1) and downstream NRF2 signaling, upregulating macropinocytosis (MP) and mitochondrial biogenesis and supporting ATP production (5, 7, 8). In contrast, intact collagen (iCol) restrains tumor growth by shutting down DDR1-NRF2 signaling through an elusive mechanism. Another poorly understood aspect of DDR1 signaling is that, despite its strong signaling capacity once activated, it exhibits unusually slow activation kinetics of its tyrosine kinase activity, distinguishing it from other receptor tyrosine kinases (RTKs) (9). Thus, while ligand availability regulates DDR1 activation, this kinetic-feature mismatch raises the possibility that additional regulatory mechanisms beyond ligand-receptor interactions exist, but these remain to be explored.

Tumor-promoting inflammation represents another defining feature of PDAC. Proinflammatory cytokines accelerate tumor initiation and progression and exert durable effects on epithelial cell behavior, consistent with the concept of inflammatory memory (10). However, how prior inflammatory exposure is molecularly encoded within tumor cells, and how this memory intersects with stromal signaling pathways to shape tumor cell state, remains poorly understood. Protein homeostasis regulated by the ubiquitin-proteasome system, and E3 ubiquitin ligases in particular, is a central determinant of tumor progression (11) and epigenetic inheritance (12, 13). Here, we identify regulated receptor degradation as a metabolic checkpoint that integrates collagen remodeling, cellular energy status, and inflammatory history to control DDR1 signaling in PDAC. We show that FBXW2, the substrate recognition subunit of an E3 ligase complex, suppresses DDR1-NRF2 signaling and is itself epigenetically silenced by cytokine-induced promoter methylation, linking inflammatory exposure to sustained alterations in receptor turnover. While providing a mechanistic basis for the “activated stroma index” characterized by increased levels of cleaved collagen (cCol) (6), our results explain how memory of past exposure to inflammatory cues is retained by the malignant cells.

## Results

### FBXW2 suppresses PDAC progression by reducing DDR1 stability

Our previous work showed that intact Col-I (iCol-I) suppresses PDAC growth, potentially by promoting DDR1 degradation (5). However, the mechanism responsible for this receptor turnover remained unknown. Consistent with post-translational control, Col-I cleavage had little effect on *DDR1* mRNA in human PDAC cell lines cultured on ECM containing either WT or MMP-resistant Col-I (R/R) (Supplementary Fig. S1A). However, the proteasome inhibitor, MG132, restored DDR1 protein levels in cells plated on R/R ECM, while the lysosomal inhibitor chloroquine had little effect (Supplementary Fig. S1B), suggesting that iCol triggers proteasome-dependent DDR1 degradation.

Proteasome degradation is a ubiquitylation-dependent process in which neddylation-dependent cullin-RING ligases (CRLs) often play key roles (11, 14). The neddylation inhibitor MLN4924 led to a dose-dependent increase in DDR1 in R/R ECM-cultured 1305 cells derived from a patient xenograft (Supplementary Fig. S1C), suggesting that DDR1 may be regulated by a cullin. To identify the relevant cullin, we immunoprecipitated (IP’ed) Flag-tagged cullins from transfected and R/R ECM-cultured 1305 cells and found DDR1 selectively associates with CUL1 (Supplementary Fig. S1D). Within CRL1 complexes, substrate recognition is mediated by F-box proteins (15). Screening major F-box proteins identified FBXW2 as the only adaptor that bound endogenous and exogenous DDR1 (Fig. 1A; Supplementary Fig. S1E and S1F), and genetic and biochemical experiments confirmed that FBXW2 controls DDR1 stability. FBXW2 ablation (FBXW2^Δ^) or knockdown (KD) increased DDR1 abundance and phosphorylation and doubled DDR1 half-life even in the absence of cleaved Col-I (cCol-I). In contrast, FBXW2^WT^ overexpression accelerated DDR1 turnover on WT ECM-plated cells (Fig. 1B; Supplementary Fig. S1G-S1J). DDR1 half-life in cells plated on WT ECM was longer than in R/R ECM-plated cells (7.5 vs. 3 h) but was shortened to around 4.5 h on FBXW2 overexpression (Supplementary Fig. S1H and S1I). Furthermore, dominant-negative, F-box deleted FBXW2 (FBXW2^ΔF^) which retains substrate recognition but cannot recruit other CRL components, increased DDR1 amounts (Supplementary Fig. S1K). Congruently, FBXW2 KD diminished DDR1 ubiquitylation, while overexpression of FBXW2^WT^, but not FBXW2^ΔF^, enhanced DDR1 ubiquitylation (Fig. 1C; Supplementary Fig. S1L). Together, these results identify CRL1^FBXW2^ as the ubiquitin ligase for iCol-I-induced DDR1 degradation, suggesting that FBXW2 may function as a molecular checkpoint limiting DDR1 signaling.

**Figure 1.**
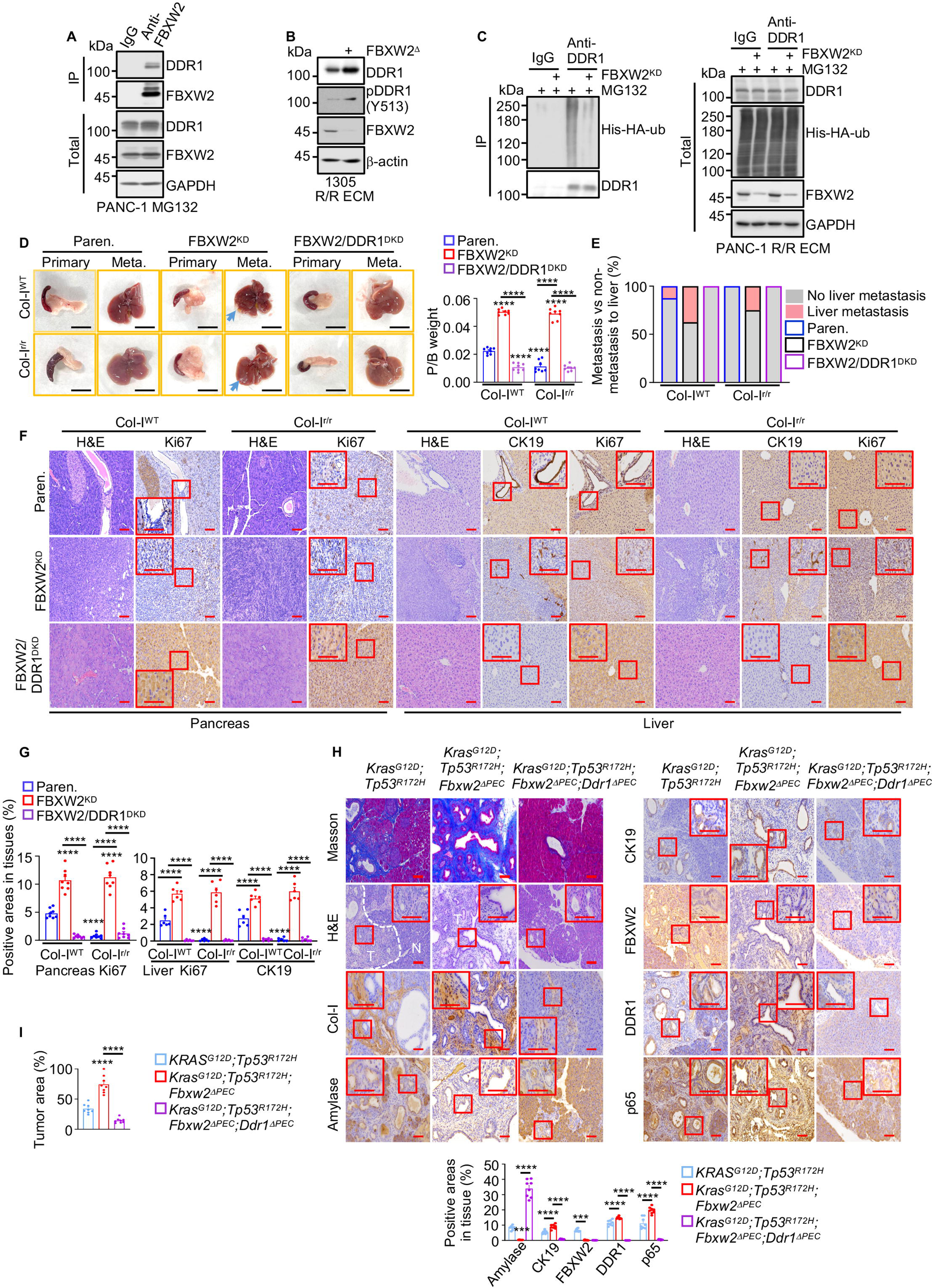
FBXW2 suppresses DDR1-dependent PDAC growth and liver metastasis. **A**, Co-IP of endogenous DDR1 with FBXW2 from PANC-1 cells treated with MG132 (5 μM) for 24h. **B**, IB analysis of the indicated proteins in parental and FBXW2^Δ^ 1305 cells grown on R/R ECM in LG medium. **C**, IB analysis of DDR1 ubiquitylation in Paren. or FBXW2 KD PANC-1 cells transfected with His-HA-ubiquitin (ub) and plated on R/R ECM in LG medium with MG132. **D**, Gross pancreas and liver morphology three weeks after orthotopic transplantation of the indicated KPC cells into Col-I^WT^ or Col-I^r/r^ mice, showing primary tumors and liver metastases (Meta.). Pancreas-to-body (P/B) weight ratio is shown to the right. **E**, Incidence of liver metastasis in mice from (**D**). **F**, Representative H&E, Ki67, and CK19 staining of pancreatic and hepatic paraffin embedded sections from (**D**). Boxed areas show higher magnification. **G**, Image J quantitation of staining intensity for the indicated proteins in (**F**). **H**, Representative Masson, H&E, and IHC staining of the indicated proteins in pancreata from the indicated mice. Quantification of positively stained areas for the indicated proteins is shown to the bottom. Pancreatic epithelial cells (PEC). **I**, Quantification of tumor areas in pancreata from (**H**) Data in (**D**, **H**, **I**) (*n*=8 mice), (**G**) (n=8 pancreatic fields or n=6 liver fields) are mean ± s.e.m. Statistical significance was determined by one-way analysis of variance (ANOVA) with Tukey post hoc tests (**D**, **G**-**I**) or Kruskal-Wallis tests with Dunn post hoc test (**H**) based on data normality distribution. Exact *P* values are shown in the Source Data. \**P* < 0.05, \*\**P* < 0.01, \*\*\**P* < 0.001, \*\*\*\**P* < 0.0001. Scale bars (**D**) 1 cm, (**F**, **H**) 100 μm.

To determine whether FBXW2-mediated DDR1 degradation impacts PDAC growth, we examined proliferation of PDAC cells with FBXW2 KD (FBXW2^KD^) or combined FBXW2 and DDR1 KD (FBXW2/DDR1^DKD^) cultured on WT or R/R ECM. As expected, R/R ECM poorly supported parental cell growth, reflecting reduced DDR1 signaling, but FBXW2 silencing restored cell proliferation and enabled efficient growth on R/R ECM (Supplementary Fig. S2A and S2B).

We next examined tumor growth in vivo. Parental, FBXW2^KD^, or FBXW2/DDR1^DKD^ KPC960 cells were orthotopically transplanted into Col-I^WT^ and Col-I^r/r^ hosts. FBXW2^KD^ cells were more tumorigenic than the parental cells and formed tumors and liver metastases in Col-I^r/r^ mice at levels comparable to the parental cells in Col-I^WT^ hosts (Fig. 1D-1G). Furthermore, regardless of Col-I cleavage, FBXW2 ablation upregulated DDR1 and its downstream signaling components, NF-κB/p65, SQSTM1/p62 and NRF2, together with the MP-related protein NHE1, and the mitochondrial protein SDHB. These effects were reversed by DDR1 ablation (Supplementary Fig. S2C). Similar results were obtained in a genetic PDAC model (*Kras^G12D^; Tp53^R172H^*, 16) in which *Fbwx2* and/or *Ddr1* was selectively ablated in pancreatic epithelial cells (PEC, Fig. 1H and 1I). These results identify FBXW2 as a key negative regulator of DDR1 whose loss enables PDAC cells to bypass the growth restrictive effects of iCol and sustain tumor progression.

### FBXW2 restrains DDR1-dependent PanIN initiation

Although we had shown that FBXW2 promotes ubiquitin-mediated DDR degradation and limits PDAC growth and metastasis, we next asked whether FBXW2 also constrains earlier stages of pancreatic tumorigenesis in vivo, during which oncogenic KRAS drives the formation of pancreatic intraepithelial neoplasia (PanIN), a principal precursor lesion of PDAC. To examine the role of FBXW2 in tumor initiation, we used *Kras^G12D/PEC^*mice which express oncogenic *Kras^G12D^* in PEC (17, 18). Low-grade preneoplastic PanIN lesions (P), composed of tall columnar cells with basally located round to oval nuclei, exhibited reduced amylase and FBXW2, and elevated Col-I, CK19, DDR1, and nuclear NF-κB/p65 compared to normal (N) acinar tissue (Fig. 2A). PEC specific ablation of *Fbxw2* in *Kras^G12D^* mice increased low-grade PanINs and acinar cell loss, indicated by decreased amylase and increased CK19 staining, while enhancing DDR1 and nuclear p65 expression, effects that were reversed in DDR1-null *Kras^G12D^;Fbxw2^ΔPEC^* mice (Fig. 2A-2C). Together with the previous results, these data indicate that a FBXW2-mediated DDR1 degradation pathway restrains PDAC development even at the earliest stages of pancreatic neoplasia.

**Figure 2.**
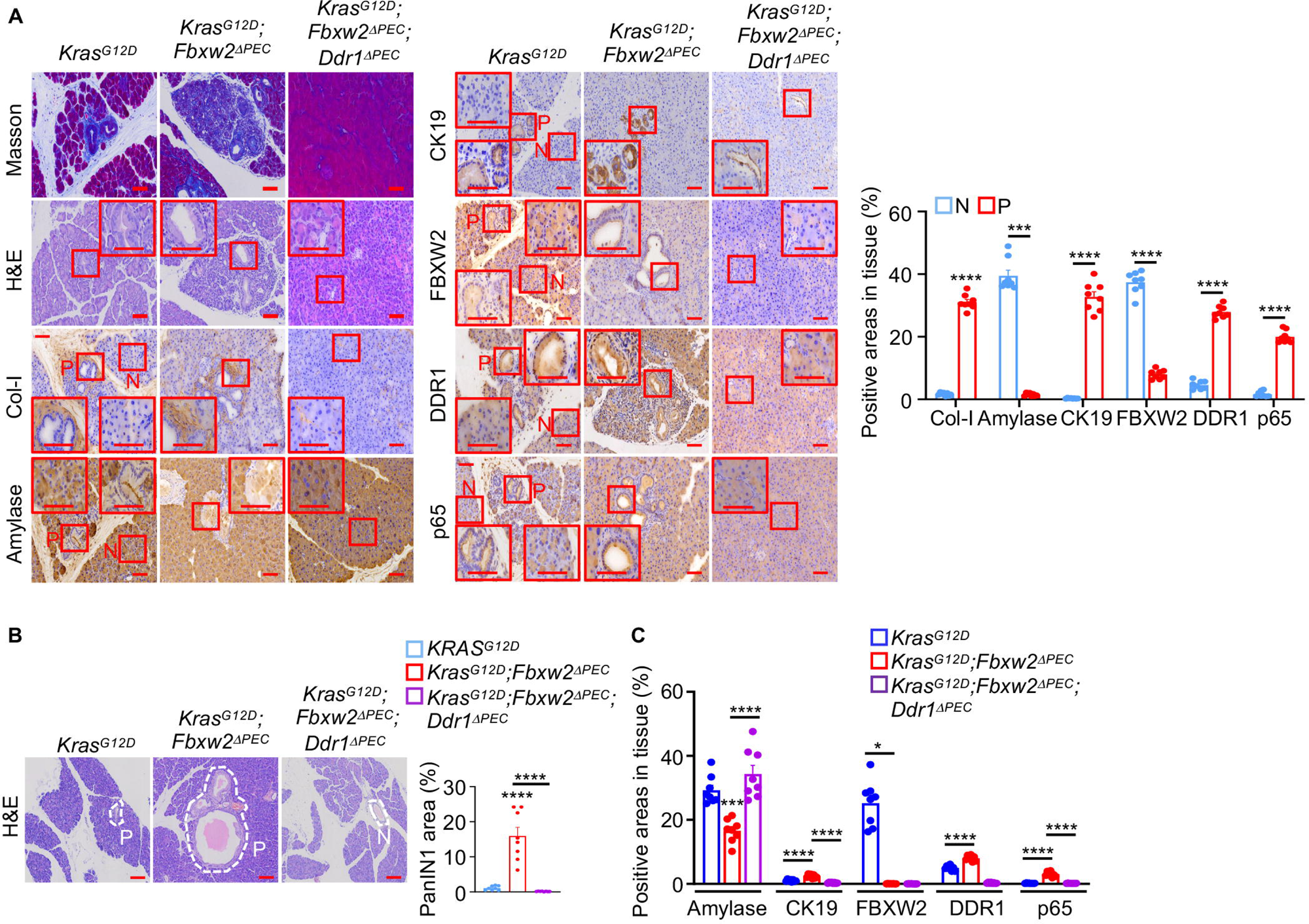
FBXW2 suppresses DDR1-dependent PanIN1 formation. **A**, Representative Masson, H&E, and IHC staining of the indicated proteins in pancreata from the indicated mice. pancreatic epithelial cells (PEC). Quantification of IHC results in normal (N) and PanIN1 (P) tissues of *Kras^G12D^* mice is shown to the right. **B**, Representative H&E staining of pancreatic sections from the indicated 8-week-old mice. Dashed circles mark PanIN1 (P) or normal ducts (N). Quantification of the PanIN1 area is shown to the right. **C**, Quantification of positively stained areas for the indicated proteins in pancreata from (**A**). Data in (**A**, **B**, **C**) (n=8 mice), are mean ± s.e.m. Statistical significance was determined by two-tailed unpaired t-test (**A**), or Mann-Whitney U-tests (**A**), one-way ANOVA with Tukey post hoc tests (**B**, **C**), or Kruskal-Wallis tests with Dunn post hoc test (**C**) based on data normality distribution. Scale bars (**A**, **B**) 100 μm.

### Intact collagen induces metabolic stress that activates AMPK-dependent DDR1 degradation

We next asked how the stromal collagen architecture translates into sustained differences in DDR1 stability. To investigate how iCol-I triggers DDR1 degradation, we generated and purified intact WT and MMP-resistant collagen (R/R) and MMP-cleaved (¼ and ¾ fragments) collagen and assessed their affinity to DDR1 by co-immunoprecipitation. ¾Col-I bound DDR1 with considerably higher affinity than intact WT or R/R collagen, whereas ¼Col-I failed to bind DDR1 (Fig. 3A). Likewise, titration experiments confirmed that iCol-I associated with DDR1 less effectively than ¾Col-I (Fig. 3B) despite the DDR1-binding motif (GVMGFO, where O denotes hydroxyproline) being present in all fibrillar collagens (19). Interestingly, AlphaFold structure prediction and protein-protein docking analyses showed that this binding motif is buried in iCol-I, and exposed in ¾Col-I (Supplementary Fig. S3A). Consistent with differential binding, PDAC cells co-cultured with iCol, ¼ Col-I or no collagen exhibited poor activation of DDR1 and its effectors (NF-κB and NRF2) and resulted in loss of DDR1 protein, whereas cells exposed to ¾Col-I maintained DDR1 abundance and signaling activity (Fig. 3C; Supplementary Fig. S3B).

**Figure 3.**
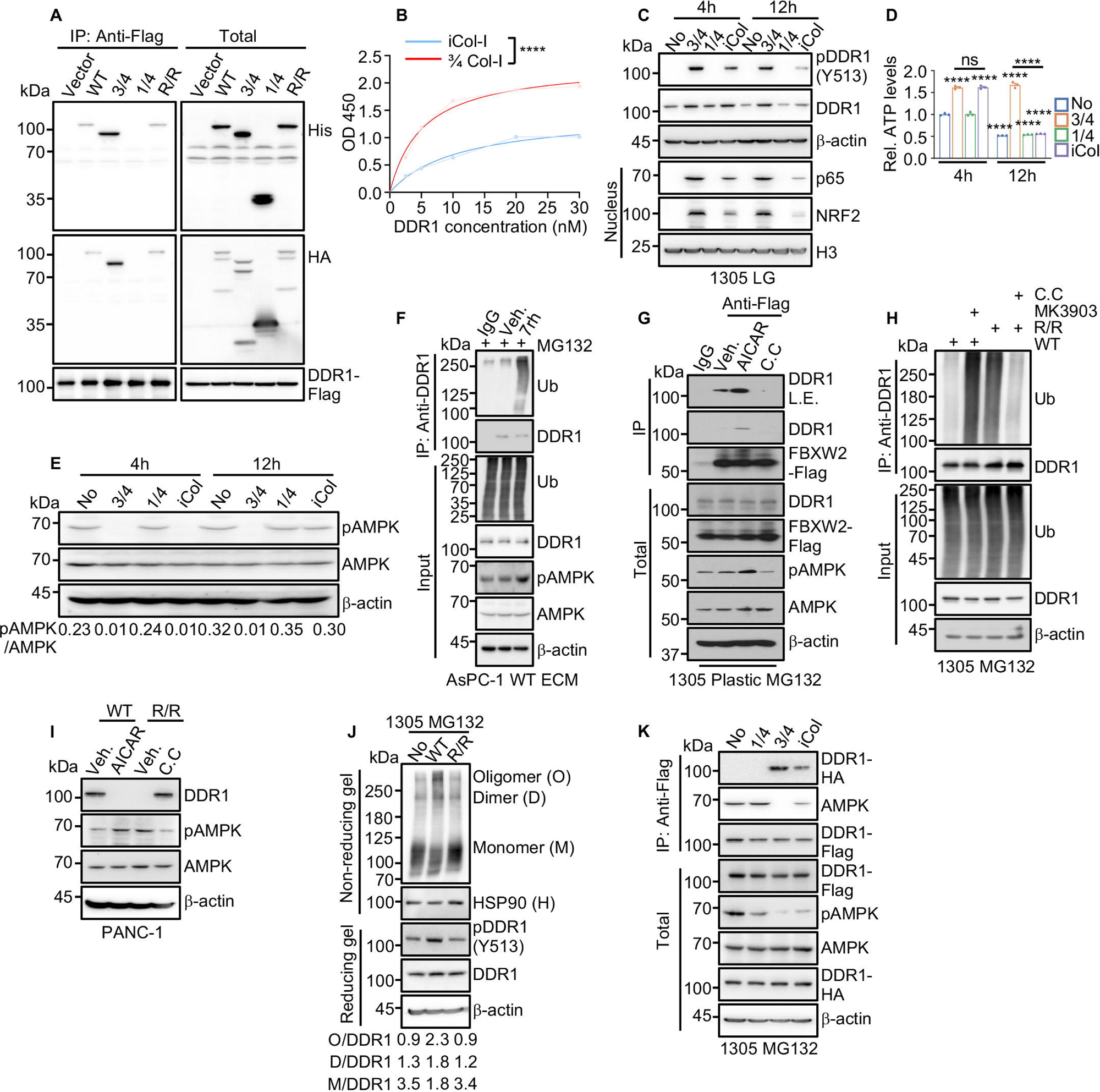
Intact Col-I induces AMPK-dependent DDR1 proteasomal degradation. **A**, Co-immunoprecipitation (co-IP) of purified DDR1-Flag with various Col-I variants and cleavage products. WT, 3/4, ¼, and R/R Col1a1 were His-tagged, and WT, ^3^/4, ¼, and R/R Col1a2 were HA-tagged. **B**, ELISA measuring binding of purified DDR1 to 3/4, or R/R Col-I (iCol-I) coated plates. Binding curves were fitted using a one-site specific binding model to determine a Kd of 5.222 nM for DDR1 and ¾ Col-I and 11.01 nM for DDR1 and iCol-I. **C**, Immunoblotting (IB) of the indicated proteins in lysates of 1305 cells treated -/+ 3/4, ¼, and intact Col-I in low-glucose (LG) medium for 4 or 12 h. **D**, Total ATP amounts in cells from (**C**), shown relative to ATP amounts in untreated (No) 1305 controls in LG medium for 4 h. **E**, IB analysis of the indicated proteins in cells from (**C**). The ratio of AMPK T172 phosphorylation (pAMPK) to total AMPK is shown below. **F**, IB analysis of DDR1 ubiquitylation in lysates of AsPC-1 cells plated on WT ECM in LG medium + MG132 (5 μM), -/+ the DDR1 inhibitor 7rh (500 nM) for 24 h. The input proteins serve as loading controls. **G**, Co-IP of DDR1 with FBXW2-Flag from 1305 cells treated with MG132 and either an AMPK activator (AICAR, 0.5 mM) or an inhibitor [Compound C (C.C), 20 μM] for 12 h. **H**, IB analysis of ubiquitylated DDR1 in 1305 cells grown on WT or R/R ECM in LG medium with MG132 and either an AMPK activator (MK3903, 20 nM) or an inhibitor (C.C, 20 μM) for 12 h. **I**, IB analysis of the indicated proteins in PANC-1 cells plated on WT or R/R ECM in LG medium -/+ AICAR or C.C. for 24 h. **J**, Non-reducing IB analysis of DDR1 isoforms from 1305 cells grown on plastic (No), WT, or R/R ECM in LG medium + MG132. The ratio of DDR1 oligomers (O), dimers (D), or monomers (M) to total DDR1 is shown below. **K**, Co-IP of DDR1-Flag with DDR1-HA and AMPK from 1305 cells treated with MG132 in LG medium in presence of the indicated Col-I forms. Data in (**B, D**) (*n*=3 independent experiments) are mean ± s.e.m. Statistical significance was determined using global fitting and an F-test (**B**) or one-way ANOVA with Tukey post hoc tests (**D**). Exact *P* values are provided in Source Data. \*\*\*\**P* < 0.0001.

By activating NRF2, cCol-I stimulates mitochondrial biogenesis and MP, thereby supporting ATP production (3, 5). Therefore, we next examined whether these differences in DDR1 signaling affected cellular energy metabolism. Indeed, after 8 h under low-nutrient conditions, intracellular ATP in cells cultured without ECM or together with ¼Col-I, iCol-I or R/R ECM was lower than in cells cultured with ¾Col-I or WT ECM (Fig. 3D; Supplementary Fig. S3C). This suggests that reduced DDR1 signaling on iCol-I imposes bioenergetic stress on tumor cells. Notably, although iCol-I reduces DDR1 activity in the short term, it does not decrease DDR1 protein levels (Fig. 3C and 3D). However, the decline in DDR1 protein coincides with a reduction in ATP levels (Supplementary Fig. S3B and S3C), suggesting that DDR1 protein stability may be linked to the cellular energy state.

Because cellular energy depletion activates AMPK, we examined AMPK activity under these conditions. The kinetics of AMPK activation in cells cultured with iCol-I or R/R ECM vs. ¾Col-I or WT ECM followed the decline in intracellular ATP with a small delay (Fig. 3E; Supplementary Fig. S3B). Consistently, inhibition of DDR1 with the inhibitor 7rh increased AMPK activity and stimulated DDR1 ubiquitylation (Fig. 3F), suggesting that metabolic stress and defective DDR1 activation leads to AMPK activation and DDR1 turnover. To specifically test whether AMPK activity regulates DDR1 stability, we treated 1305 cells with the AMPK activator AICAR. Activation of AMPK enhanced FBXW2 binding to DDR1, while the AMPK inhibitor, compound C (C.C) blunted this interaction (Fig. 3G). The AMPK activator, MK3903, promoted DDR1 ubiquitylation in WT ECM-cultured 1305 cells, which was blunted by C.C. (Fig. 3H). Consistent with this, AMPK activation resulted in reduced DDR1 protein levels, and AMPK inhibition reversed this in R/R ECM-cultured cells (Fig. 3I). Together, these results suggest a model in which iCol reduces DDR1 signaling and cellular ATP levels, thereby activating AMPK to promote ubiquitin-mediated DDR1 degradation.

DDR1 activation relies on lateral dimerization and subsequent oligomerization which enhances binding to immobilized Col-I and facilitates inter-dimer phosphorylation (20, 21). Non-reducing gel electrophoresis revealed that WT, but not R/R, ECM dramatically promoted DDR1 dimerization, oligomerization, and autophosphorylation (Fig. 3J). IP analysis showed that ¾Col-I and WT ECM enhanced DDR1 self-association and inhibited AMPK-DDR1 interactions (Fig. 3K; Supplementary Fig. S3D). Conversely, the total absence of ECM, or incubation with ¼Col-I, iCol-I or R/R ECM blocked DDR1 self-association and promoted its binding to AMPK (Fig. 3K; Supplementary Fig. S3D). These results suggest that iCol-I induces DDR1 degradation because its inferior DDR1 binding activity compromises downstream NRF2 signaling and ATP production, resulting in AMPK activation. Thus, cellular adenylate charge defines a DDR1 stability regime, rather than an acute on-off response.

### DDR1 proteostasis controls metabolic fitness and tumor growth

Our prior results indicate that metabolic stress induced by iCol activates AMPK and promotes DDR1 degradation. We next sought to determine how AMPK signaling couples cellular energy status DDR1 proteostasis. We therefore tested whether DDR1 is a direct substrate of AMPK. *In vitro* kinase assay demonstrated that recombinant HA-DDR1 was directly phosphorylated by AMPK (Fig. 4A). Using protein mass spectrometry (MS), we identified T519 in the DDR1 cytoplasmic region as a potential AMPK phosphoacceptor (Supplementary Fig. S3E). Replacement of T519 with alanine (A) and transfection of 1305 cells cultured on R/R ECM with WT and T519A DDR1 HA-tagged expression vectors validated T519 as the AMPK phosphoacceptor (Fig. 4B). As expected, the T519A substitution prevented R/R ECM or AMPK induced DDR1 degradation (Fig. 4C; Supplementary Fig. S3F).

**Figure 4.**
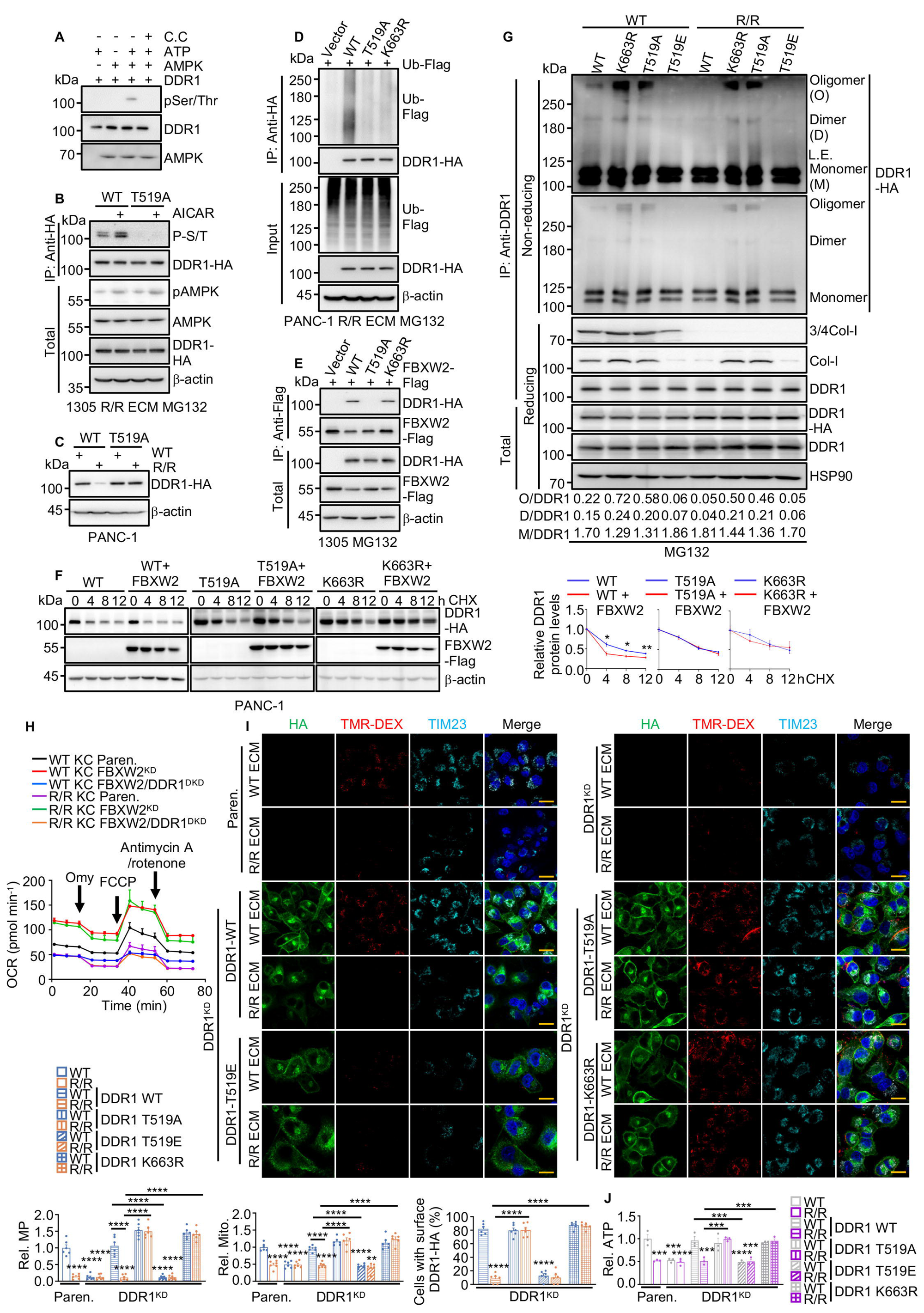
DDR1 phosphorylation by AMPK promotes its ubiquitylation by FBXW2. **A**, *In vitro* kinase assay with purified active AMPK and Flag-tagged DDR1 isolated from 293T cells. **B**, IB analysis of WT or T519A DDR1-HA from1305 cells cultured on R/R ECM in LG medium + MG132, -/+ AICAR treatment for 24 h. **C**, IB analysis of the indicated proteins in WT or T519A DDR1-HA-expressing PANC-1 cells cultured on WT or R/R ECM in LG medium. **D**, IB analysis of IP’ed DDR1-HA variants co-expressed with ubiquitin-Flag in PANC-1 cells grown on R/R ECM in LG medium + MG132. **E**, IB analysis of FBXW2-Flag co-IP’ed with the indicated DDR1-HA variants from transfected 1305 cells grown in LG medium + MG132. **F**, IB analysis of DDR1-HA variants co-expressed -/+ FBXW2-Flag in PANC-1 cells treated with 100 μg/ml cycloheximide (CHX) for the indicated durations. The graph shows quantification of relative DDR1 amounts. **G**, DDR1-HA variants and the indicated proteins were co-IP’ed from 1305 cells grown on WT or R/R ECM in LG medium + MG132 and separated on reducing and non-reducing gels and IB analyzed. The ratio of DDR1 oligomers (O), dimers (D), or monomers (M) to IP’ed total DDR1 (reducing gel) is shown below. Degradation-resistant DDR1 variants (K663R and T519A) show increased oligomerization compared to WT DDR1. **H**, Oxygen consumption rate (OCR) of parental, FBXW2^KD^, and FBXW2/DDR1^DKD^ KC cells plated on WT or R/R ECM and incubated in LG medium for 24 h before and after treatment with oligomycin (Omy), FCCP, or rotenone/antimycin A. **I**, Representative images and quantification of mitochondria (TIM23), macropinocytosis (MP, TMR-dextran uptake), and surface DDR1 in parental and DDR1^KD^ PANC-1 cells reconstituted with the indicated DDR1 variants and cultured on WT or R/R ECM in LG medium. Scale bar, 20 μm. **J**, Total ATP amounts in cells from (**I**), normalized to parental PANC-1 cells plated on WT ECM. Data in (**F**, **H**, **I**) (*n*=3 independent experiments) and (**I**) (*n*=6 fields) are presented as mean ± s.e.m. Statistical significance was assessed using two-tailed unpaired t-tests (**F**), or one-way ANOVA with Tukey post hoc tests (**I**, **J**). Exact *P* values are provided in Source Data. \**P* < 0.05, \*\**P* < 0.01, \*\*\**P* < 0.001, \*\*\*\**P* < 0.0001.

MS analysis of DDR1 from R/R-ECM cultured cells identified K663 as a potential ubiquitylation site (Supplementary Fig. S3G). Like T519A, the K663R substitution abrogated DDR1 ubiquitylation, but importantly, only the T519A mutation impaired FBXW2 recruitment (Fig. 4D and 4E). Both substitutions extended DDR1 half-life to 12 h, even in FBXW2-overexpressing cells (Fig. 4F). The phosphomimic T519E mutation had the opposite effect, blunting DDR1 oligomerization, interaction with collagen, and tyrosine phosphorylation, and shortened its half-life from 8 to 4 h in WT ECM-cultured cells (Fig. 4G; Supplementary Fig. S3H and S3I). The K663R and T519A mutations also enhanced DDR1 dimerization and oligomerization and binding to ¾ and full-length Col-I and autophosphorylation (Fig. 4G; Supplementary Fig. S3H).

Notably, FBXW2 KD increased rates of oxygen consumption, mitochondrial abundance, and MP activity, regardless of the Col-I status, a DDR1 dependent effect (Fig. 4H, Supplementary Fig. S3J). Re-expression of the T519A and K663R variants, which exhibit prominent cell surface localization, in DDR1 KD cells plated on either WT or R/R ECM restored mitochondrial number, ATP content, and MP activity, metabolic processes that support PDAC growth in nutrient-limited environments, while unmodified DDR1 restored these parameters only in WT ECM-cultured cells (Fig. 4I and 4J). Re-expression of the phosphomimic T519E mutant, which is predominantly cytoplasmic, didn’t rescue any of these phenotypes (Fig. 4I and 4J). Accordingly, cells expressing the T519A or K663R DDR1 variants were refractory to suppression of PDAC growth, liver metastasis, and NF-κB and NRF2 activities by iCol-I (Supplementary Fig. S3K-S3O). Together, these results identify a metabolic checkpoint in which AMPK phosphorylation of DDR1 at T519 promotes FBXW2-dependent ubiquitination and degradation, linking cellular energy status to receptor stability and metabolic adaptation.

### Low FBXW2 and reduced DDR1 T519 phosphorylation predict worse clinical outcome

To assess whether the AMPK-DDR1-FBXW2 axis reflects a stable signaling state in human disease, we analyzed surgically resected PDAC specimens by immunohistochemistry (IHC). The majority of tumors (65/93) exhibited high levels of cleaved collagen (cCol-I^hi^) and low expression of FBXW2 (FBXW2^lo^, 55/93). Notably, most FBXW2^lo^ tumors exhibited increased staining for DDR1 and downstream signaling components including p65, NRF2, SDC1, and SDHB, consistent with activation of the DDR1-driven program (Fig. 5A and 5B; Supplementary Fig. S4A-S4C). Among these markers, only DDR1 and p65 exhibited statistically significant inverse correlations with FBXW2, supporting a functional link between FBXW2 loss and DDR1 pathway activation (Fig. 5A and 5B; Supplementary Fig. S4A and S4C). In addition, we found that DDR1 T519 phosphorylation (pT519-DDR1) was inversely correlated with DDR1 and NRF2 staining (Fig. 5A and 5B). Accordingly, most DDR1^hi^ tumors were pT519^lo^ (44/59) (Fig. 5B). These findings suggest that FBXW2 loss in human PDAC enables DDR1 activation by cCol-I. Moreover, pT519-DDR1 inversely correlated with the activating DDR1 phosphorylation at Y513 (Fig. 5A and 5B).

**Figure 5.**
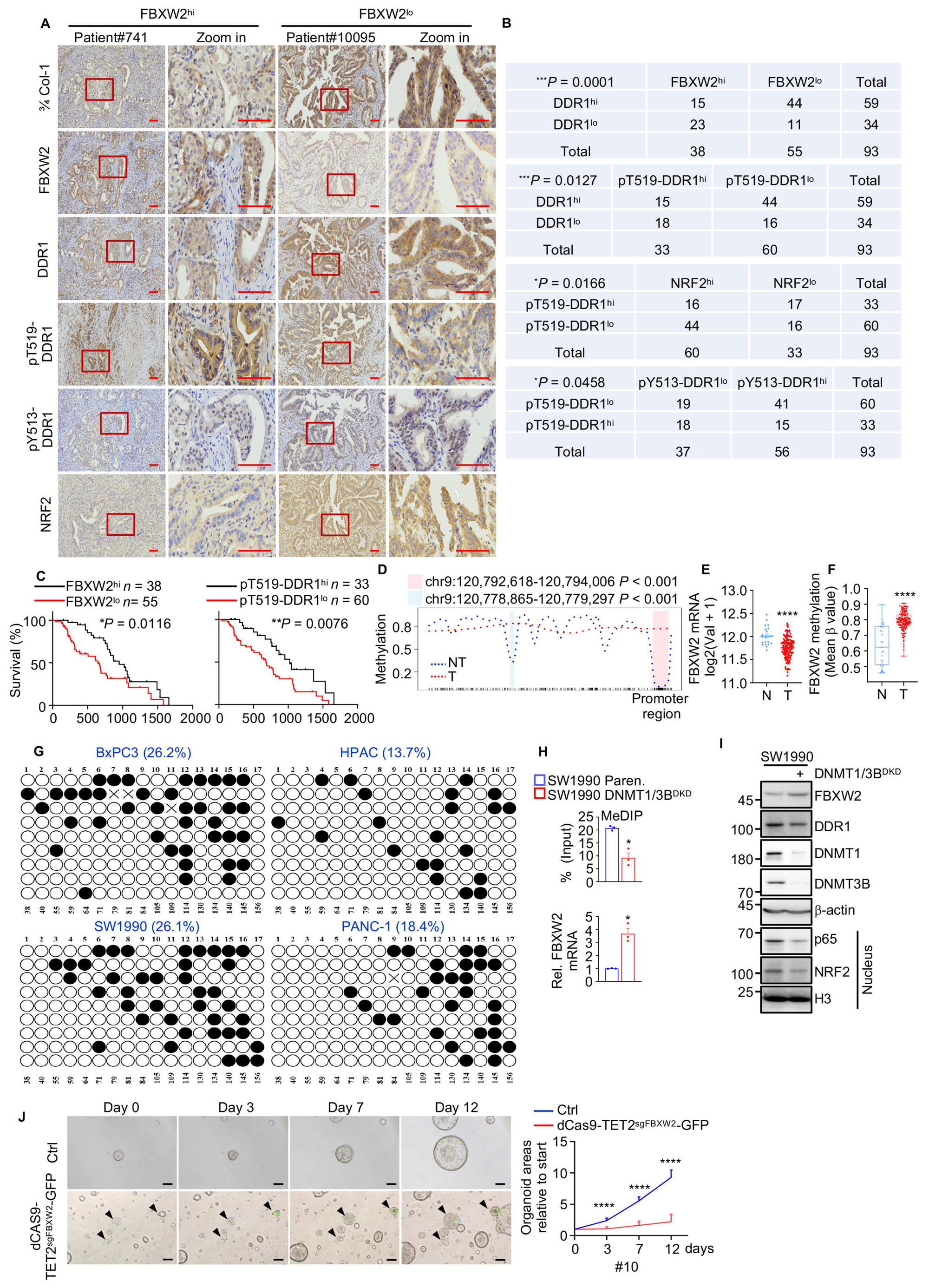
Low FBXW2 expression correlates with poor clinical outcome in human PDAC. **A**, Representative IHC of resected FBXW2^hi^ (#741) and FBXW2^lo^ (#10095) human PDAC tissues. Boxed areas are further magnified. **B**, Correlation between expression levels of the indicated proteins in human PDAC specimens examined by a two-tailed Chi-square test. \**P* < 0.05, \*\*\**P* < 0.001. **C**, Comparisons of overall survival between patients with PDAC stratified according to FBXW2 or pT519-DDR1. Significance was determined by log-rank test. \**P* < 0.05, \*\**P* < 0.01. **D**, Methylation frequency at individual CpG sites in the *FBXW2* promoter and 5’ region [Chr. 9: 120,771,978-120,793,438 (extended=1000)] in human PDAC tumor (T, *n*=7) and tumor-adjacent (N, *n*=3) tissues. **E**, *FBXW2* mRNA amounts in human PDAC tumor (T, *n*=140) and tumor-adjacent (N, *n*=21) tissues, based on data from the CPTAC-PDAC database. **F**, FBXW2 promoter DNA methylation levels in human PDAC tumor (*n*=167) and tumor-adjacent (*n*=29) tissues, based on data from GSE49149. **G**, *FBXW2* DNA promoter region methylation determined by bisulfite sequencing (BS)-PCR analysis of the indicated cell lines. **H**, Methylated DNA immunoprecipitation (MeDIP)-qPCR analysis of *FBXW2* promoter methylation in Paren. and DNMT1/3B^DKD^ FBXW2^lo^ (SW1990) cells. *FBXW2* mRNA levels in the same cells were determined by qPCR. **I**, IB analysis of the indicated proteins in cells from (**H**). **J**, Representative images of human PDAC organoids (#10) -/+ overexpressing TET2 targeting to the *FBXW2* promoter (dCas9-TET2^sgFBXW2^-GFP). Total organoids areas were measured by Image J. Data are presented relative to each individual organoid on day 0 and shown to the right. Arrows indicate organoids with dCas9-TET2^sgFBXW2^-GFP. Data in (**H**) (*n*=3 independent experiments) and (**J**) (*n*=15 organoids) are presented as mean ± s.e.m. Statistical significance was assessed using two-tailed unpaired t-tests (**E**, **H**), or Mann-Whitney test (**F**, **J**) based on data normality distribution. Exact *P* values are shown in the Source Data. \**P* < 0.05, \*\**P* < 0.01, \*\*\**P* < 0.001, \*\*\*\**P* < 0.0001. Scale bars (**A**) 100 μm, (**J**) 50 μm.

Analysis of the Clinical Proteomic Tumor Analysis Consortium database for PDAC (CPTAC-PDAC) showed that pT519-DDR1 was elevated in early-stage (I-II) PDAC but low in advanced disease (stage III-IV), whereas pY513-DDR1 showed the opposite pattern (Supplementary Fig. S4D). Phosphosite compatibility analysis showed that pT519 was mutually exclusive with activating tyrosine phosphorylations (pY513, pY792, or pY796), further indicating that pT519 marks inactive, degradation-destined DDR1 (Supplementary Fig. S4E). Consistent with these observations, patients with FBXW2^hi^ or pT519-DDR1^hi^ tumors had longer median survival than those with the opposite profile (Fig. 5C). These observations raise the possibility that FBXW2 loss reflects an actively programmed, durable tumor state, rather than a passive byproduct of tumor evolution.

### *FBXW2* is epigenetically silenced by promoter hypermethylation in human PDAC

The durability of FBXW2 loss in human PDAC suggested that it may encode a form of epigenetic memory rather than transient regulation. Consistent with this, *FBXW2* mutations are rare in human PDAC, suggesting that epigenetic mechanisms may suppress its expression. scATAC-seq revealed reduced chromatin accessibility at the *FBXW2* promoter region (chr9:120,792,300-120,792,700) in malignant epithelial cells compared to normal epithelial cells, correlating with lower *FBXW2* mRNA expression (Supplementary Fig. S5A-S5C). Moreover, FBXW2^hi^ tumors (Pri2) exhibited greater promoter accessibility than FBXW2^lo^ tumors (Pri3) (Supplementary Fig. S5D). MethMarkerDB-PDAC dataset analysis showed that the *FBXW2* promoter was hypermethylated in tumors relative to normal tissues (Fig. 5D). This result was validated in data obtained from the public cohorts: decreased tumoral *FBXW2* mRNA was accompanied by promoter hypermethylation compared to adjacent non-tumor tissues (Fig. 5E and 5F). *FBXW2* promoter methylation was validated by bisulfite sequencing (BS-seq) of human PDAC cell lines stratified according to relative *FBXW2* expression (Fig. 5G; Supplementary Fig. S5E). *FBXW2* promoter methylation was elevated in FBXW2^lo^ PDAC cells (BxPC3 and SW1990) who had high DDR1, relative to FBXW2^hi^ cell lines (HPAC and PANC-1) (Fig. 5G; Supplementary Fig. S5E). These findings indicate that DNA methylation inversely correlates with FBXW2 expression and DDR1 signaling activity.

To determine whether promotor methylation directly regulates FBXW2 expression, we inhibited DNA methyltransferases DNMT1 and DNMT3A/B by genetic ablation or with 5-azacytidine (5-AzC). In FBXW2^lo^ cells and organoids, demethylation reduced *FBXW2* promoter methylation, restored FBXW2 expression and reduced DDR1 expression and signaling (Fig. 5H and 5I; Supplementary Fig. S5F-S5H). Conversely, overexpression (OE) DNMT1 and DNMT3B in FBXW2^hi^ cells increased *FBXW2* promoter methylation, reducing its expression, and increasing DDR1 expression and signaling, whereas overexpression of TET2 methylcytosine dioxygenase had the opposite effect (Supplementary Fig. S5I-S5K). In addition, FBXW2^lo^ cells were more sensitive to 5-AzC, consistent with dependence on DNA methylation for maintenance of this state (Supplementary Fig. S5L). Finally, TET2 targeting to the *FBXW2* promoter using a dCas9-TET2 fusion guided by *FBXW2* promoter-recognizing sgRNA (dCas9-TET2^sgFBXW2^-GFP) slowed the growth of human PDAC organoids (Fig. 5J).

### Cytokine-induced *FBXW2* promoter methylation

Inflammatory signaling not only activates NF-kB and STAT3 transcriptional programs (22), but also can promote epigenetic remodeling (23). Given that memory of past exposure to inflammatory cues has been shown to stimulate emergence and malignant progression of low-grade PanIN lesions (10), we examined whether inflammatory cytokines induced *FBXW2* promoter methylation. IL-17A and IL-8 or CXCL1/2, its murine homologs (24), markedly decreased FBXW2 mRNA and protein and upregulated DDR1 expression. Importantly, ectopic expression FBXW2 reversed DDR1 upregulation, indicating that cytokine-induced DDR1 activation is mediated through FBXW2 suppression (Fig. 6A-6C; Supplementary Fig. S6A). BS-seq and MeDIP-qPCR showed that IL-8 and IL-17A induced *FBXW2* promoter methylation in FBXW2^hi^ human PDAC cells and organoids, and this was reversed by DNMT1 and DNMT3B depletion or inhibition (Fig. 6D and 6E; Supplementary Fig. S6B and S6C). Congruently, IL-17A and IL-8 silenced FBXW2 expression and increased DDR1 and nuclear p65 and NRF2 in FBXW2^hi^ PDAC organoids and cell lines cultured in LG medium on either WT or R/R ECM (Fig. 6F and 6G; Supplementary Fig. S6D). These changes were abolished by DNMT inhibition or dCas9-TET2^sgFBXW2^ targeting (Fig. 6F and 6G; Supplementary Fig. S6D). Notably, IL-8 suppressed R/R ECM-induced pT519-DDR1 amounts and enhanced MP activity and mitochondrial content, effects that were reversed by dCas9-TET2^sgFBXW2^ or DNMT1/3B depletion (Fig. 6G; Supplementary Fig. S6E). Unlike WT DDR1, the T519A variant was stably expressed in cells cultured on R/R ECM with LG medium, regardless of the IL-8- or dCas9-TET2^sgFBXW2^-induced changes in FBXW2 expression (Supplementary Fig. S6F).

**Figure 6.**
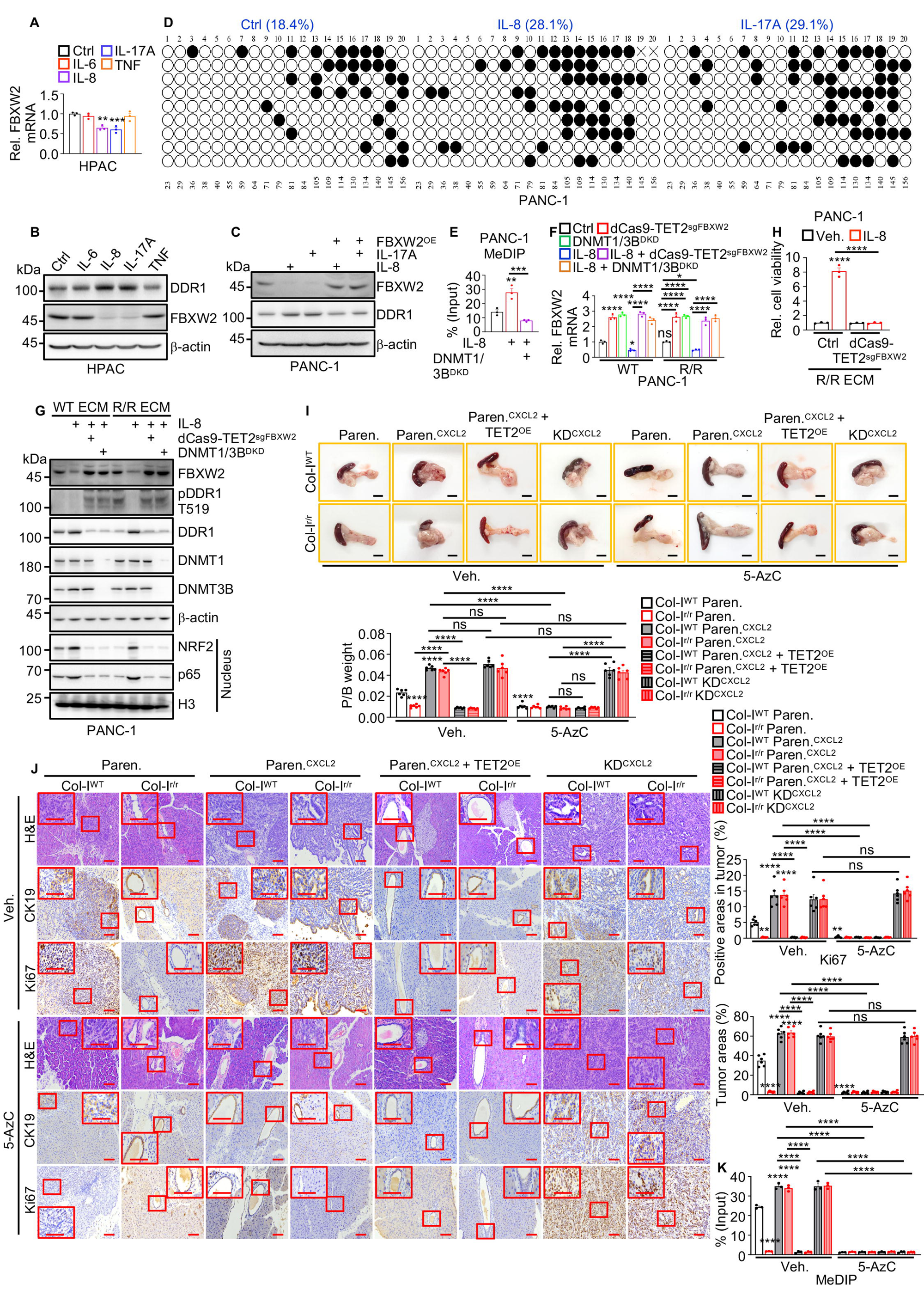
Inflammatory cytokines stimulate PDAC growth by inducing FBXW2 promoter methylation. **A**, qPCR analysis of *FBXW2* mRNA in HPAC cells treated -/+ 20 ng/ml IL-6, IL-8, IL-17A, or TNF for 7 days. **B**, IB analysis of the indicated proteins in cells from (**A**). **C**, IB analysis of the indicated proteins in Paren. and FBXW2^OE^ PANC-1 cells treated -/+ IL-8 or IL-17A for 7 days. **D**, *FBXW2* DNA promoter region methylation validated by BS-PCR analysis of PANC-1 cells treated -/+ IL-8 or IL-17A for 7 days. **E**, MeDIP-qPCR analysis of *FBXW2* promoter methylation in Paren. and DNMT1/3B^DKD^ PANC-1 cells treated -/+ IL-8 for 5 days. **F**, qPCR analysis of *FBXW2* mRNA in PANC-1 cells treated -/+ IL-8 for 5 days following DNMT1/3B^DKD^ or TET2 targeting to the *FBXW2* promoter (dCas9-TET2^sgFBXW2^), and subsequently grown on WT or R/R ECM in LG medium for 24 h. **G**, IB analysis of the indicated proteins in cells from (**F**). **H**, PANC-1 cells were pretreated -/+ IL-8 for 5 days, following TET2 targeting to *FBXW2*, and then grown on R/R ECM in LG medium for 3 days after which cell viability was assessed and normalized to that of untreated cells. **I**, Gross pancreatic morphology and P/B weight ratio 3 weeks after orthotopic transplantation of Paren., Fbxw2-KD (KD) KC cells pretreated -/+ CXCL2 for 5 days, followed by -/+ TET2 targeting to the *Fbxw2* promoter (TET2^OE^) into Col-I^WT^ or Col-I^r/r^ mice -/+ 5-AzC treatment initiated on day 3 post-transplantation. **J**, Representative H&E and IHC of indicated proteins in pancreata from (**i**). Boxed areas are further magnified. Quantification of tumor Ki67 staining and tumor areas is shown on the right. **K**, MeDIP-qPCR analysis of *Fbxw2* promoter methylation in EpCAM^+^ tumor cells isolated from pancreata in (**I**). Data in (**A**, **E**, **F**, **H**, **K**) (*n*=3 independent experiments) and (**I**, **J**) (*n*=6 mice) presented as mean ± s.e.m. Statistical significance was assessed using one-way ANOVA with Tukey post hoc tests. Exact *P* values are shown in the Source Data. \**P* < 0.05, \*\**P* < 0.01, \*\*\**P* < 0.001, \*\*\*\**P* < 0.0001. Scale bars (**I**) 1 cm, (**J**) 100 μm.

### Inflammatory priming establishes a durable epigenetic memory that sustains DDR1 signaling and tumor fitness independent of collagen state

We next asked whether cytokine-induced FBXW2 silencing represents a transient response or establishes a durable memory of inflammatory exposure that persists after cytokine withdrawal. PDAC cells pretreated with IL-8 or CXCL-2 were cultured under nutrient-limited conditions on either WT or R/R ECM, or orthotopically transplanted into Col-I^WT^ and Col-I^r/r^ mice. IL-8/CXCL-2 pretreatment accelerated PANC-1, HPAC and KC cell growth irrespective of Col-I cleavage. This effect was abolished by targeted demethylation of the FBXW2 promoter using dCas9-TET2^sgFBXW2^ or DNMT3B ablation, and restored by FBXW2 depletion (Fig. 6H; Supplementary Fig. S6). Similar methylation dependent effects were observed with IL-17A and IL-8 in human PDAC organoids, and were reversed by 5-AzC (Supplementary Fig. S6H). Accordingly, CXCL2 pretreatment promoted KC tumor growth and liver metastases, upregulated Ki67, DDR1, p65, and NRF2, and suppressed FBXW2 expression relative to untreated cells in both Col-I^WT^ and Col-I^r/r^ hosts (Fig. 6I and 6J3; Supplementary Fig. S7A-S7D). Moreover, CXCL2-pretreated PDAC cells displayed enhanced tumor growth, Ki67, FBXW2, DDR1, p65, and NRF2 expression, and liver metastases in Col-I^r/r^ mice like cells in Col-I^WT^ hosts. Conversely, targeted demethylation of the *FBXW2* promoter abrogated these effects, whereas 5-AzC treatment had a similar effect on WT, but not FBXW2-depleted cells (Fig. 6I and 6J; Supplementary Fig. S7A-S7D). Of note, prior exposure to IL-8/CXCL2 or IL-17 led to a sustained upregulation of DNMT3B and increased *FBXW2* promoter methylation both in vitro, and tumors derived from pretreated cells (Fig. 6K; Supplementary Fig. S7E and S7F). These effects were independent of Col-I cleavage, suggesting that inflammatory cues induce long-lasting DNMT3B-mediated FBXW2 silencing that overrides microenvironment constraints. Together, these results show that inflammatory priming establishes an epigenetic memory that silences FBXW2, preserves DDR1 signaling, and sustains tumor growth independent of stromal context (Fig. 7).

**Figure 7.**
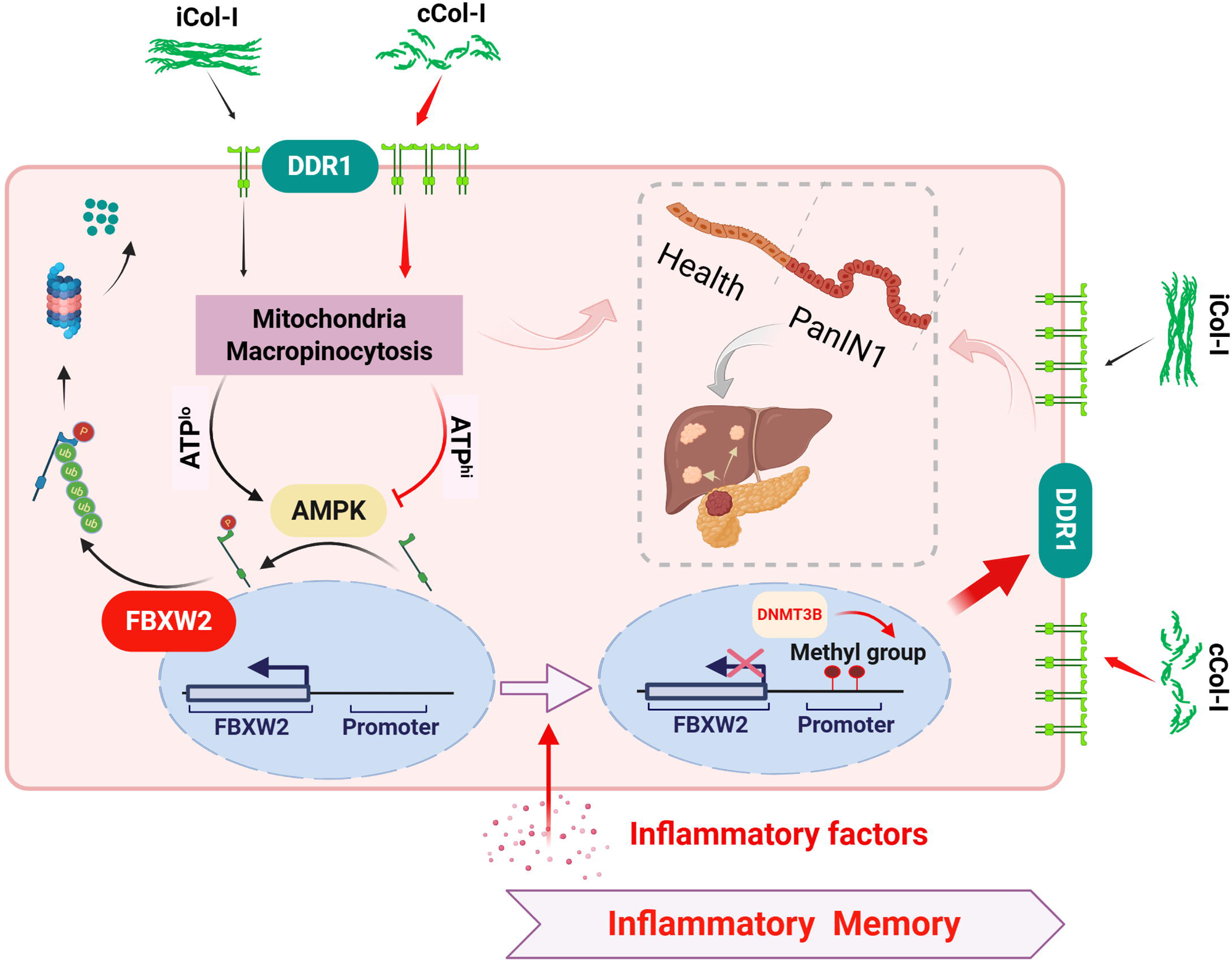
A metabolic-inflammatory checkpoint governs DDR1 stability and tumor progression. Impaired Col-I remodeling limits the high-affinity DDR1 ligand (¾Col-I), reducing ATP levels and activating AMPK, which drives DDR1 phosphorylation and FBXW2-dependent degradation. Inflammatory signaling overrides this pathway by inducing DNMT3B-mediated silencing of FBXW2, establishing an inflammatory memory that stabilizes DDR1 and sustains tumor progression even under stromal constraint.

## Discussion

Tumor progression within PDAC is shaped by the interaction between cancer cells and the collagen-rich extracellular matrix. While our previous work demonstrated that MMP generated collagen fragments activate DDR1 signaling to promote tumor growth (3, 5), the present study reveals an unanticipated layer of regulation. Here, we show that DDR1 is not only activated by the microenvironment, but it is actively destabilized in response to metabolic stress, and this degradation itself is subject to epigenetic control, dependent on inflammatory history (Fig. 7).

Although our earlier work postulated that iCol-I downregulates DDR1 by inducing an alternative receptor conformation (25), here we identify a metabolic-proteostatic checkpoint in which iCol-I, by failing to efficiently activate DDR1, leads to reduced mitochondrial output, MP activity, and a decline in cellular ATP (Fig. 7). Specifically, we found that iCol-I induces DDR1 degradation due to its poor DDR1 binding, thereby precluding activation of NRF2, the key DDR1 effector that stimulates mitochondrial biogenesis and ATP production (5, 8, 18, 26). Thus, iCol-I causes a slow decline in the adenylate charge, energetic stress, and consequent activation of AMPK. Surprisingly, AMPK regulates DDR1 itself, by phosphorylates residue T519, licensing its recognition by the E3 ligase adaptor FBXW2 and promoting proteasomal degradation. In this manner, extracellular matrix architecture is translated into receptor stability through cellular energy sensing. Importantly, this unorthodox mechanism of RTK downregulation also applies to human PDAC, where elevated pT519-DDR1 correlates with diminished DDR1 activation, marked by pY513-DDR1, and improved patient survival, indicating that this pathway operates in clinically relevant disease states.

Our data also provides detailed insight into how collagen processing regulates DDR1 activation. The GVMGFO motif mediating DDR1 binding is present in fibrillar collagens (19), but the relative affinity of different Col-I cleavage forms to DDR1 was not explored. AlphaFold modelling shows that the hydrophobic GVMGFO motif is only partially exposed in iCol-Iα1 but is made accessible after MMP-mediated cleavage, explaining the weak activity of iCol-I. Formation of high order DDR1-¾Col-I aggregates causes persistent DDR1 activation and downstream signaling required for PDAC survival in the nutrient poor desmoplastic microenvironment. Of note, DDR1 activation promotes MMP expression (27), establishing a vicious cycle that supports PDAC metabolic adaptation and rapid growth. The latter is maintained by sustained NRF2 activation, which stimulates mitochondrial biogenesis (5) and upregulates the epigenetic modifier EZH2 (18). Although previous studies suggest that DDR1 is constitutively dimerized (28), our findings indicate that DDR1 dimerization and oligomerization, which protect it from degradation, are tightly controlled by MMP-mediated Col-I cleavage. In this respect, DDR1 with its slow and sustained activation kinetics (29) is radically different from other RTKs that undergo rapid ligand induced activation followed by internalization, monoubiquitylation and degradation (30).

Our findings further establish FBXW2 as a critical tumor suppressor in PDAC, whose primary function in this context is to constrain DDR1 signaling. FBXW2 was described as a suppressor of lung, prostate, and breast cancers due to its ability to induce β-catenin (31), EGFR (32), and moesin (33) degradation. Curiously, however, FBXW2 was also shown to enhance lung and breast cancer growth and contribute to chemotherapy resistance by degrading MSX2 (34). However, except for the rare loss-of-function S84C mutation (35), the mechanisms accounting for FBXW2 loss in cancer were heretofore unknown. Here, we demonstrate that loss of FBXW2 is not primarily driven by genetic alteration, but by epigenetic silencing, and that this loss is a determinant of malignant progression and metastatic spread, with its tumor-suppressive activity largely mediated through control of DDR1 stability.

Our results also explain how memory of past exposure to tumor promoting cytokines is retained by PDAC cells and translated into sustained alterations in receptor proteostasis and signaling competence. We show that exposure to tumor-promoting cytokines including IL-17A or IL-8 silences *FBXW2* transcription through promoter methylation, thereby enhancing activation of DDR1 by ¾Col-I and subsequent tumor growth. Although promoter hypermethylation has been implicated in loss of several tumor suppressors (36), including the HCC suppressor FBP1 (37), induction of DNA methylation by tumor promoting cytokines was previously unknown.

Functionally, this epigenetic memory decouples tumor behavior from stromal constraints. Cytokine-primed cells exhibited sustained DDR activation, enhanced mitochondrial capacity and increased tumor growth, regardless of collagen cleavage status. Thus, whereas collagen architecture normally governs DDR1 stability through metabolic stress, inflammatory priming overrides this regulation by locking cells into a DDR1 active states. These findings provide a mechanistic explanation for how transient inflammatory exposures can have long lasting effects on tumor progression.

---

**Figure S1.**
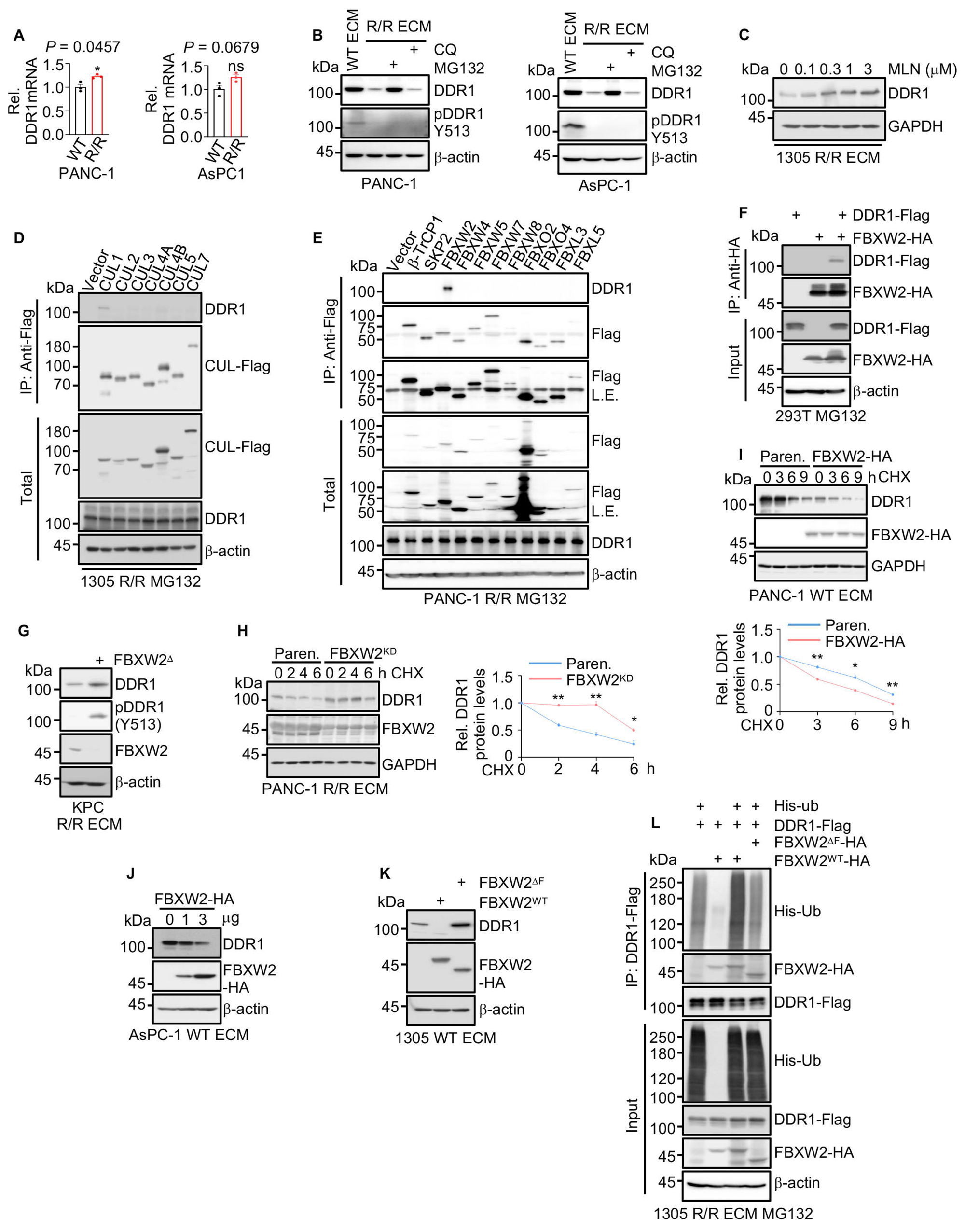
The E3 ligase CUL1^FBXW2^ mediates intact Col-I-induced DDR1 ubiquitylation and proteasomal degradation. **A**, qPCR analysis of *DDR1* mRNA in PANC-1 and AsPC-1 cells cultured on WT or R/R ECM in low glucose (LG) medium for 24 h. **B**, IB analysis of the indicated proteins in PANC-1 and AsPC-1 cells cultured as above -/+ the lysosomal inhibitor chloroquine (CQ, 50 μM) or the proteasome inhibitor MG132 (5 μM). **C**, IB analysis of DDR1 in 1305 cells cultured on R/R ECM in LG medium with the indicated concentrations of the neddylation inhibitor MLN4924 for 24 h. **D**, Co-IP of endogenous DDR1 with the Flag-tagged cullins from 1305 cells cultured on R/R ECM in LG medium with MG132 for 24 h. **E**, Co-IP of endogenous DDR1 with Flag-tagged F-box proteins from PANC-1 cells treated as above. **F**, Co-IP of DDR1-Flag with FBXW2-HA from MG132-treated (24h) 293T cells. **G**, IB analysis of the indicated proteins in parental and FBXW2^Δ^ KPC960 (KPC) cells grown on R/R ECM in LG medium. **H**, **I**, IB analysis of DDR1 in parental (Paren.) and FBXW2 knock-down (KD) PANC-1 cells plated on R/R ECM (**H**), and in Paren. and FBXW2-HA overexpressing (OE) PANC-1 cells plated on WT ECM (**I**) in LG medium and treated with 100 μg/ml CHX for the indicated durations. The graphs show relative DDR1 amounts. **J**, IB analysis of endogenous DDR1 in AsPC-1 cells transfected with increasing amounts of FBXW2-HA expression vector and cultured on WT ECM in LG medium. **K**, IB analysis of endogenous DDR1 in 1305 cells expressing either HA-tagged WT FBXW2 (FBXW2^WT^) or F-box-deleted FBXW2 (FBXW2^ΔF^), cultured as in (**J**). **L**, Co-IP of FBXW2-HA or FBXW2^ΔF^-HA with DDR1-Flag and IB analysis of ubiquitylated DDR1-Flag in 1305 cells co-transfected -/+ His-ub and plated on R/R ECM in LG medium with MG132. Data in (**A**, **H**, **I**) (n=3 independent experiments) are presented as mean ± s.e.m. Statistical significance was assessed using two-tailed unpaired t-tests. Exact *P* values are provided in the Source Data. \**P* < 0.05, \*\**P* < 0.01.

**Figure S2.**
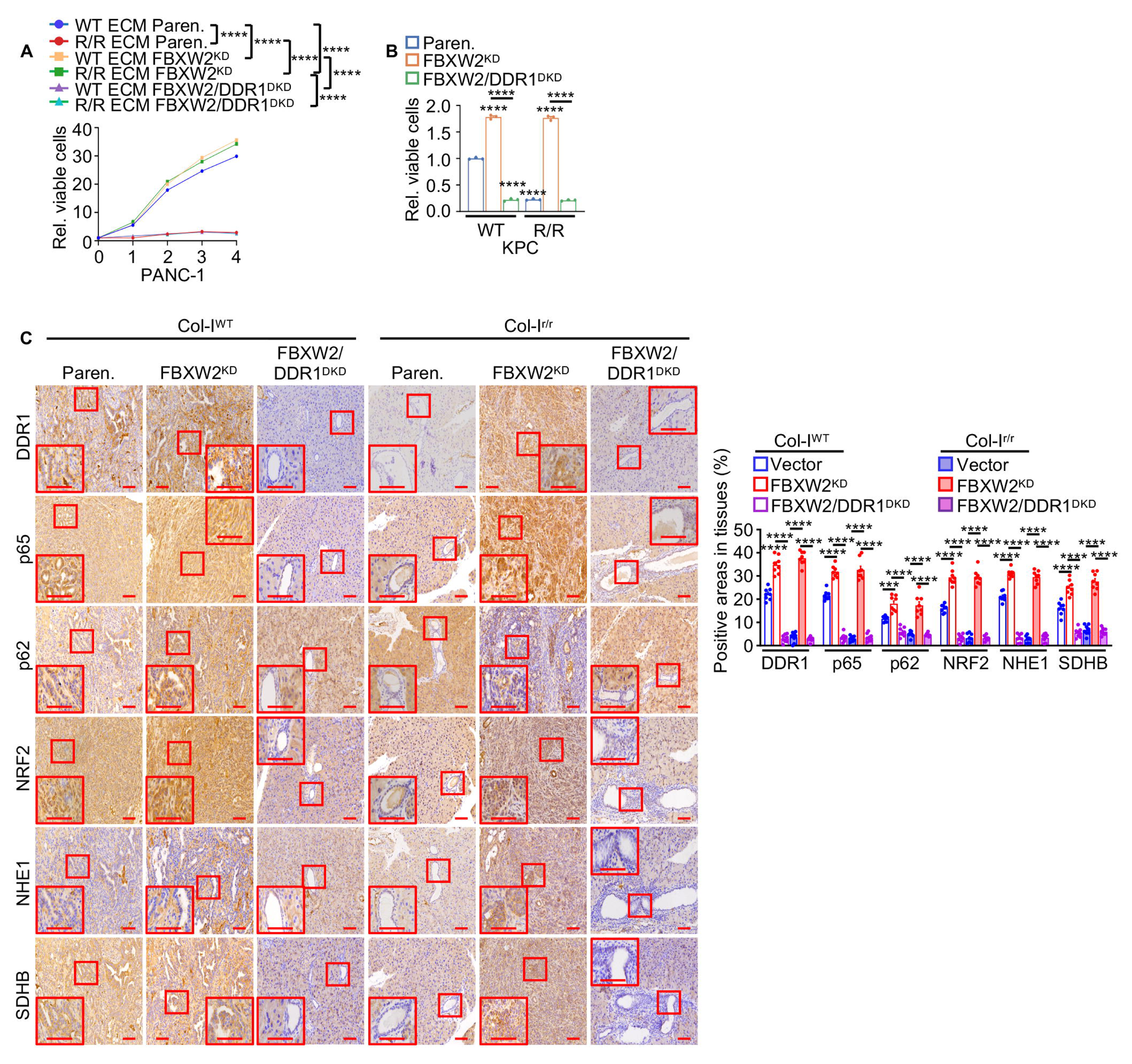
FBXW2 deficiency promotes PDAC tumor growth in a DDR1-dependent manner. **A**, Paren., FBXW2^KD^, and FBXW2/DDR1^DKD^ PANC-1 cells were cultured on WT or R/R ECM in LG medium. Viable cells were measured at the indicated timepoints, and their numbers are shown relative to day 0. **B**, Viability of the indicated KPC960 (KPC) cells treated as in (**A**) measured after 3 days in LG medium and normalized to that of Paren. cells plated on WT ECM. **C**, Representative IHC of indicated proteins in pancreata three weeks after orthotopic transplantation of the indicated KPC cells into Col-I^WT^ or Col-I^r/r^ mice. Boxed areas are further magnified. Quantification of IHC results is shown to the right. Results in (**A**, **B**) (*n*=3 independent experiments), and (**C**) (*n*=8 mice) are mean ± s.e.m. Statistical significance was determined using one-way ANOVA with Tukey post hoc tests (**A**-**C**). Exact *P* values are provided in the Source Data. \*\*\**P* < 0.001, \*\*\*\**P* < 0.0001. Scale bars (**C**) 100 μm.

**Figure S3.**
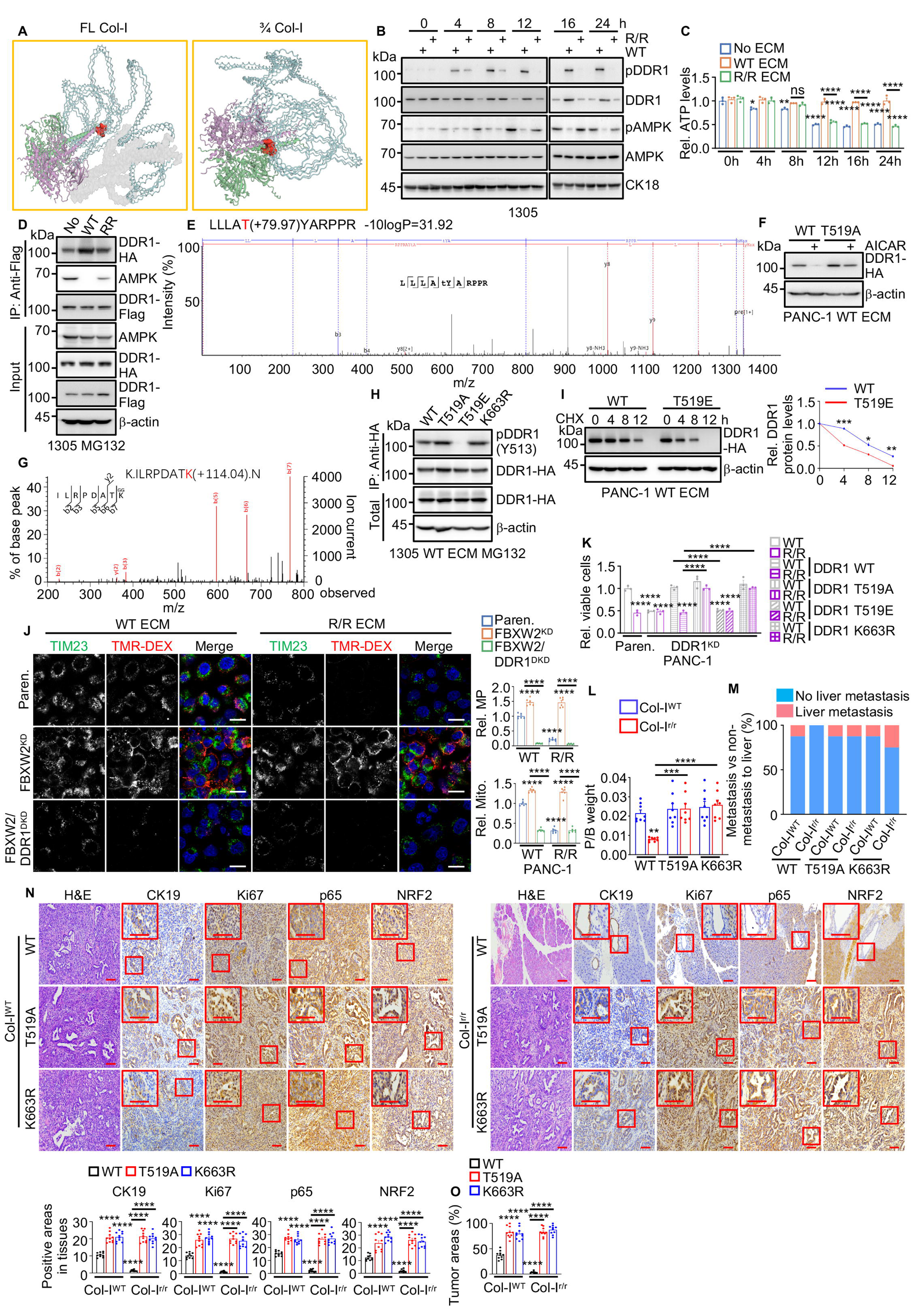
iCol-I-induced AMPK activation facilitates DDR1 degradation. **A**, AlphaFold3 structural prediction coupled with HDock analysis showing the interaction between full-length (FL) Col-I (docking score: -234.64, left) or ¾Col-I (docking score: -285.55, right) and DDR1. The DDR1-binding motif (GVMGFO) is highlighted in red. The N-terminus of FL or ¾Col-I is shown in blue, the C-terminus in grey, and the DDR1 dimer in green and pink. ¾Col-I displays stronger and more stable binding to DDR1. **B**, IB analysis of 1305 cells cultured on plastic, WT ECM, or R/R ECM in LG medium for the indicated durations after cell attachment. **C**, Total cellular ATP in above cells is presented relative to its amount in cells grown on plastic at the 0 h timepoint. **D**, Co-IP of DDR1-HA and AMPK with DDR1-Flag from 1305 cells cultured on plastic, WT, or R/R ECM in LG medium with MG132 for 24 h. **E**, Mass spectrometry identification of the AMPK phosphorylation site of DDR1. Shown is the MS/MS spectrum of a tryptic peptide with a +79.97 Da shift indicative of a phosphate group at the T residue. **F**, IB analysis of HA-tagged DDR1 variants expressed in PANC-1 cells cultured on WT ECM in LG medium -/+ AICAR for 24 h. **G**, MS identification of the DDR1 ubiquitylation site. Shown is the MS/MS spectrum of a tryptic peptide with a +114.04 Da shift indicative of a ubiquitin group at the K residue. **H**, IB analysis of DDR1 activation (Y513 phosphorylation) in 1305 cells expressing different DDR1-HA variants, cultured on WT ECM in LG medium + MG132. **I**, IB analysis of WT and T519E DDR1 in PANC-1 cells plated on WT ECM in LG medium and treated with 100 μg/ml CHX for the indicated durations. The graphs show relative DDR1 amounts. **J**, Representative images and quantification of mitochondria (TIM23) and macropinocytosis (TMR-DEX uptake) in Paren., FBXW2^KD^, and FBXW2/DDR1^DKD^ PANC-1 cells cultured on WT or R/R ECM in LG medium for 24 h. **K**, Paren. and DDR1^KD^ PANC-1 cells that either express or not the indicated DDR1 variants were plated on WT or R/R ECM in LG medium. Cell viability was assessed after three days and normalized to that of the parental cells plated on WT ECM. **L**, P/B weight ratio three weeks after orthotopic transplantation of KC cells expressing the indicated DDR1 variants into Col-I^WT^ or Col-I^r/r^ mice. **M**, Incidence of liver metastasis in mice from (**L**). **N**, Representative H&E, CK19, Ki67, p65, and NRF2 staining of pancreatic sections from Col-I^WT^ and Col-I^r/r^ mice 3 weeks after orthotopic transplantation of KC6141 (KC) cells expressing WT, T519A, or K663R DDR1. Boxed areas show higher magnification. Quantification of positively stained areas for the indicated proteins is shown to the right. **O**, Quantification of tumor areas in mouse pancreata from (**N**). Data in (**C**, **I**, **K**) (*n*=3 independent experiments), (**J**) (*n*=6 fields), (**L**) (*n*=8 mice), and (**N**, **O**) (*n*=10 fields) are presented as mean ± s.e.m. Statistical significance was assessed using one-way ANOVA with Tukey post hoc tests (**C**, **J**-**L**, **N**, **O**) or two-tailed unpaired t-tests (**I**). Exact *P* values are provided in the Source Data. \**P* < 0.05, \*\**P* < 0.01, \*\*\**P* < 0.001, \*\*\*\**P* < 0.0001. Scale bars (**J**) 20 μm, (**N**) 100 μm.

**Figure S4.**
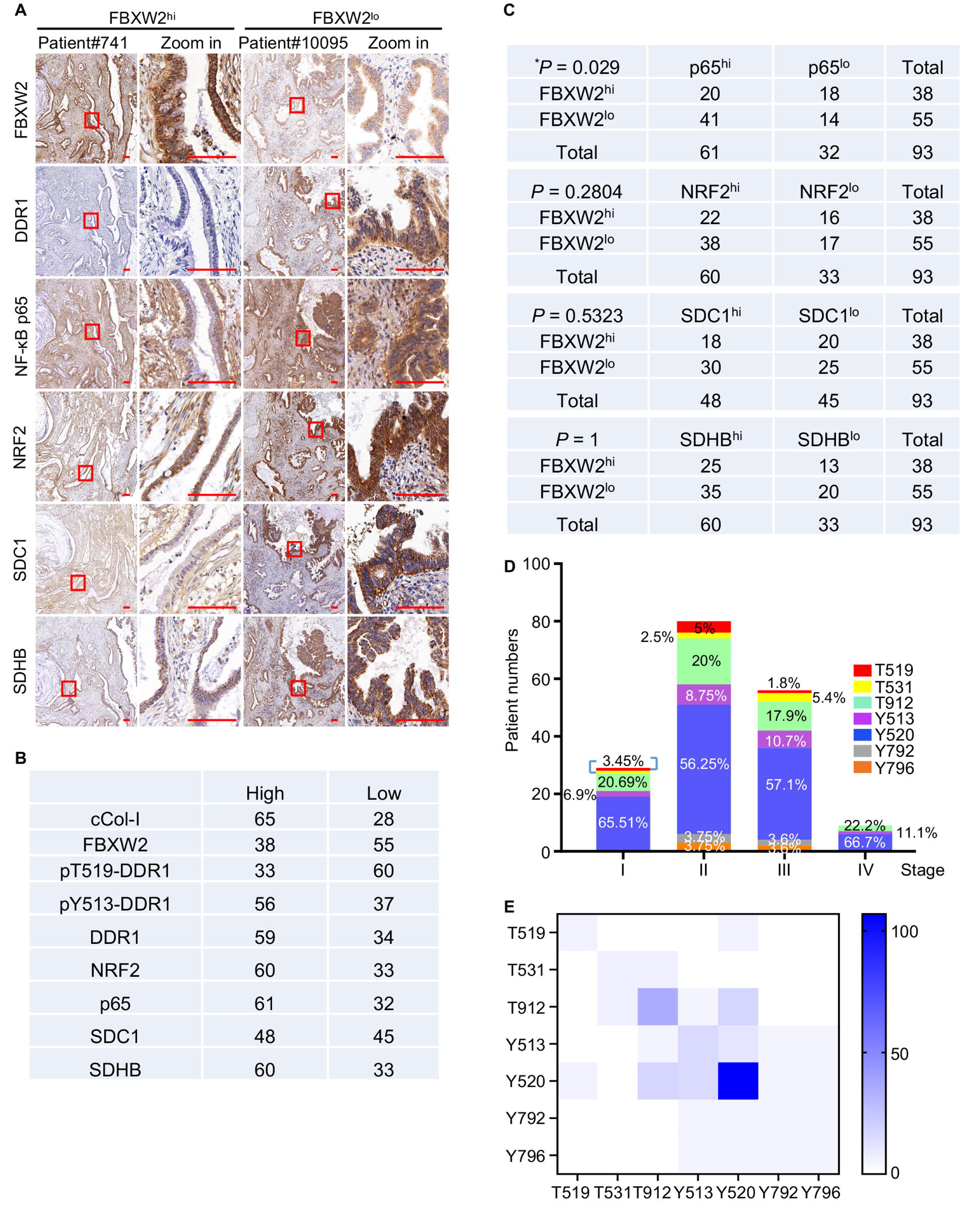
The FBXW2-DDR1 axis in human PDAC specimens. **A**, Representative IHC of resected human PDAC tissues. Boxed areas are further magnified. Scale bars, 100 μm. **B**, Numbers of human PDAC specimens (*n*=93) positive for the indicated proteins, indicated as low and high expression. **C**, Correlation between the indicated proteins in the above specimens was analyzed by a two-tailed Chi-square test. \**P* < 0.05. **D**, Distribution of different DDR1 phosphorylation states in stage I, II, III, and IV PDAC specimens. DDR1 phosphorylation data were procured from the CPTAC-PDAC database. **E**, Phosphosite compatibility analysis of the indicated DDR1 phosphorylation sites in human PDAC using data derived from CPTAC-PDAC.

**Figure S5.**
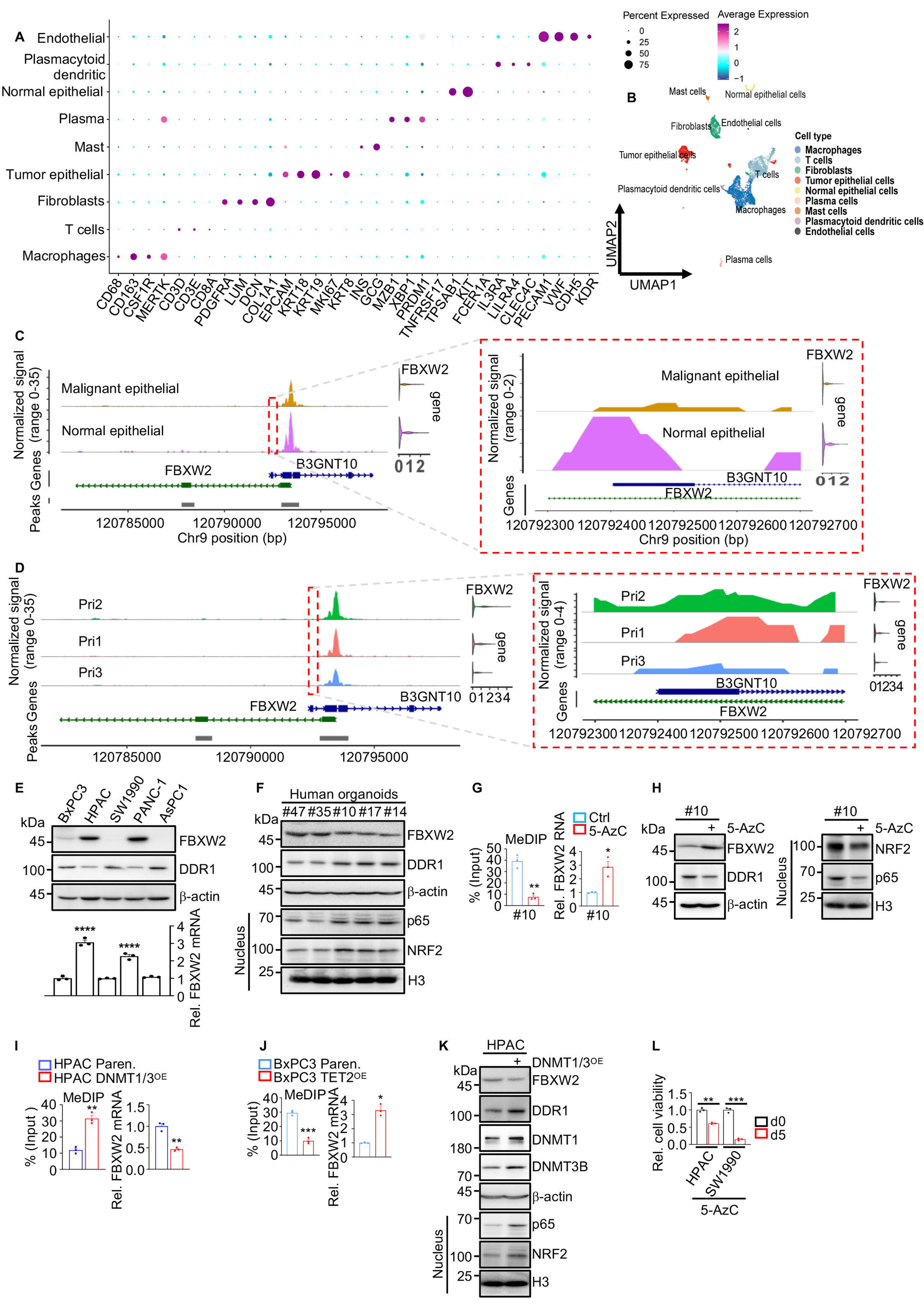
*FBXW2* promoter hypermethylation in pancreatic tumors suppresses its transcription. **A**, Dotplots showing the percentages of the indicated cell types and average expression levels of canonical marker genes across 9 cell clusters identified from the single nuclear assay for transposase accessible chromatin (ATAC) sequencing (snATAC-Seq) of human PDAC [HTAN DCC Portal (https://data.humantumoratlas.org/), Sample ID: HTA12_24_4]. **B**, UMAP visualizations and unsupervised clustering of 6,058 nuclei from the above dataset. **C**, ATAC-seq tracks of the *FBXW2* locus in normal and malignant epithelial cells, with differentially accessible promoter regions (Chr9:120,792,300-120,792,700) highlighted by a red dashed box. Relative *FBXW2* mRNA amounts are shown to the right. **D**, ATAC-seq tracks of the *FBXW2* locus in PDAC malignant epithelial cells from three patients (HTAN DCC Portal, Pri1, Pri2, and Pri3). Differentially accessible promoter regions (Chr9:120,792,300-120,792,700) are highlighted by a red dashed box. Higher promoter accessibility correlated with increased *FBXW2* mRNA expression. **E**, IB and qPCR analyses of the indicated proteins in human PDAC cell lines, showing an inverse correlation between FBXW2 and DDR1 expression levels. **F**, IB analysis of the indicated proteins in human PDAC organoids, showing an inverse correlation between FBXW2 expression and DDR1-NRF2 axis activity. **G**, MeDIP-qPCR analysis of *FBXW2* promoter methylation in a FBXW2^lo^ human PDAC organoid (#10) treated -/+ 5-AzC for three weeks. Relative *FBXW2* mRNA amounts are shown to the right. **H**, IB analysis of the indicated proteins in human PDAC organoids from (**G**). **I**, **J**, MeDIP-qPCR of *FBXW2* promoter methylation in Paren. and DNMT1/3B^OE^ FBXW2^hi^ (HPAC) cells (**I**) or in FBXW2^lo^ human PDAC cells (BxPC3) -/+ TET2^OE^ (**J**). *FBXW2* mRNA amounts in the same cells were determined by qPCR. **K**, IB analysis of the indicated proteins in cells from (**I**). **L**, HPAC and SW1990 cells were plated on WT ECM in LG medium + 5-AzC for 5 days. Cell viability was assessed at day 0 (d0) and day 5 (d5) and normalized to d0. Data in (**E**, **G**, **I**, **J**, **L**) (*n*=3 independent experiments) are mean ± s.e.m. Statistical significance was assessed by one-way ANOVA with Tukey post hoc tests (**E**, **L**) or two-tailed unpaired t-tests (**G**, **I**, **J**). Exact *P* values are shown in the Source Data. \**P* < 0.05, \*\**P* < 0.01, \*\*\**P* < 0.001.

**Figure S6.**
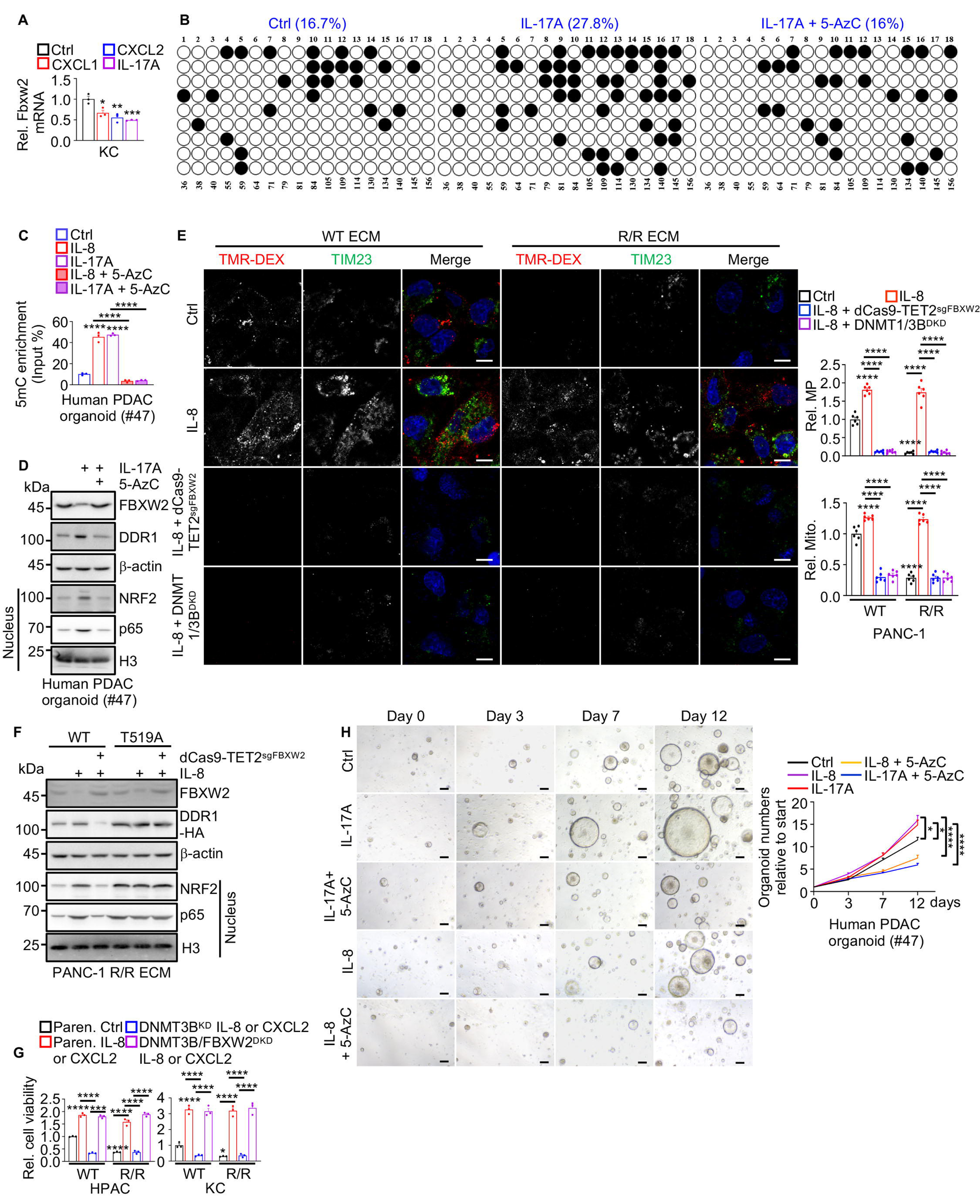
Inflammatory cytokines induce *FBXW2* promoter hypermethylation, resulting in reduced FBXW2 expression and enhanced DDR1-NRF2 axis activity. **A**, qPCR quantitation of *Fbxw2* mRNA in mouse PDAC KC6141 (KC) cells -/+ the indicated inflammatory cytokines for 5 days. **B**, **C**, *FBXW2* DNA promoter region methylation determined by BS-seq (**B**) or MeDIP-qPCR (**C**) in a FBXW2^hi^ human PDAC organoid (#47) -/+ IL-8, IL-17A, or 5-AzC for 3 weeks. **D**, IB analysis of the indicated proteins in a human PDAC organoid (#47) treated -/+ IL-17A, or 5-AzC for 3 weeks. **E**, Representative images and quantification of MP (TMR-DEX uptake) and mitochondria (TIM23) in PANC-1 cells pretreated -/+ IL-8 for 5 days, following TET2 targeting to *FBXW2* promoter or DNMT1/3B^DKD^, and subsequently grown on WT or R/R ECM in LG medium for 24 h. **F**, IB analysis of the indicated proteins in WT and T519A DDR1 OE PANC-1 cells treated -/+ IL-8 for 5 days, following TET2 targeting to the *FBXW2* promoter and subsequently grown on R/R ECM in LG medium for 24 h. **G**, Paren., DNMT3B^KD^, and DNMT3B/FBXW2^DKD^ HPAC or KC cells pretreated -/+ IL8 (HPAC) or CXCL2 (KC) for 5 days, then grown on WT or R/R ECM in LG medium -/+ IL-8 or CXCL2 for 3 days after which cell viability was determined and normalized to that of untreated Paren. cells plated on WT ECM. **H**, Representative images of human PDAC organoids (#47) treated -/+ IL-8, IL-17A, or 5-AzC as indicated. Total organoids areas were measured by Image J. Data are presented relative to each individual organoid on day 0 and shown on the right. Data in (**A**, **C**, **G**) (*n*=3 independent experiments), (**E**) (*n*=6 fields), and (**H**) (*n*=27 organoids) are mean ± s.e.m. Statistical significance was assessed by one-way ANOVA with Tukey post hoc tests. Exact *P* values are shown in the Source Data. \**P* < 0.05, \*\**P* < 0.01, \*\*\**P* < 0.001, \*\*\*\**P* < 0.0001. Scale bars (**E**) 10 μm, (**H**) 50 μm.

**Figure S7.**
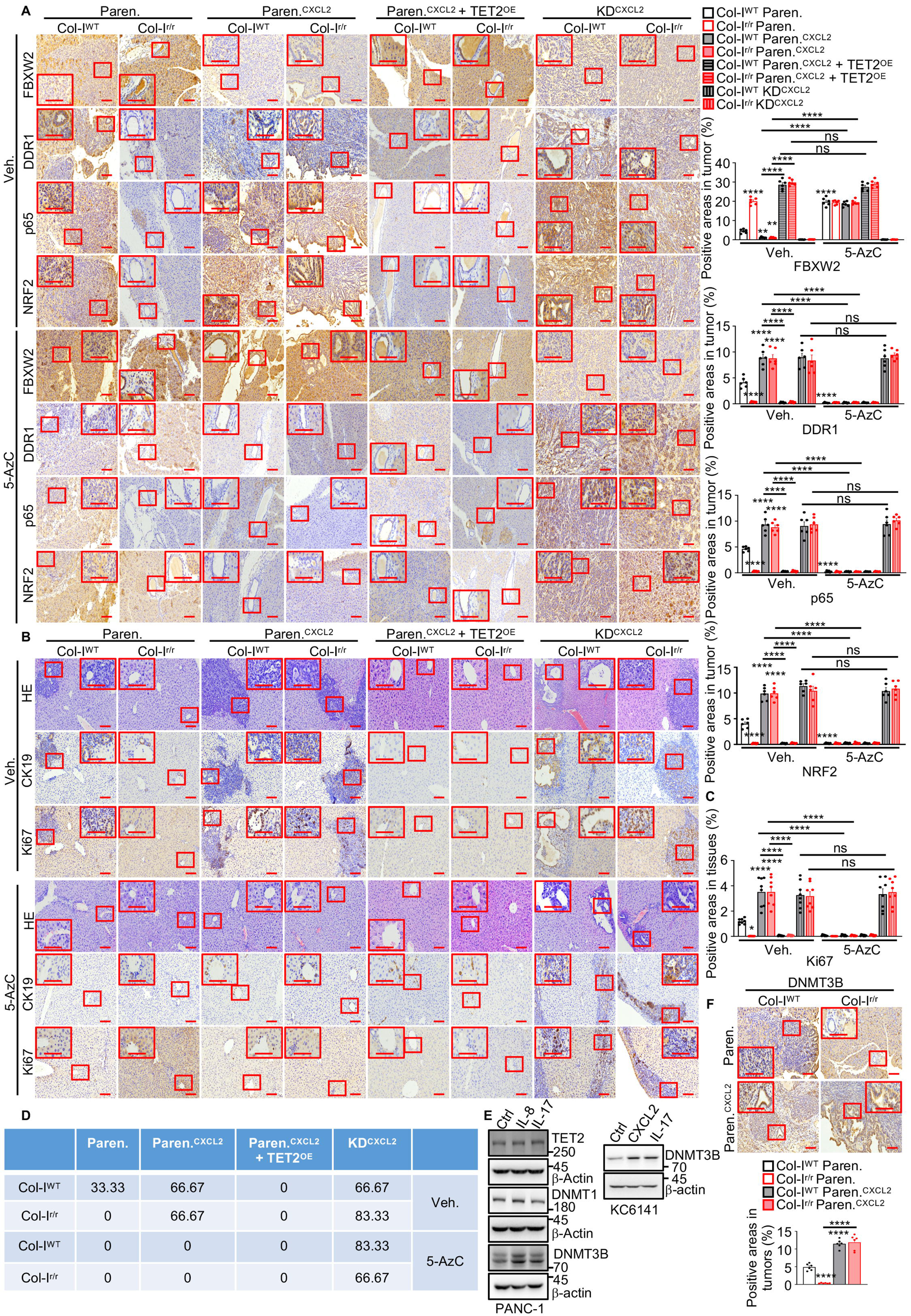
Inflammatory cytokines promote PDAC growth and liver metastasis by inducing DNMT3B expression and *FBXW2* promoter methylation. **A**, Representative IHC of indicated proteins in pancreata three weeks after orthotopic transplantation of Paren., Fbxw2-KD (KD) KC cells pretreated -/+ CXCL2 for 5 days, followed by -/+ TET2 targeting to the *Fbxw2* promoter (TET2^OE^) into Col-I^WT^ or Col-I^r/r^ mice -/+ 5-AzC treatment initiated on day 3 post-transplantation. Boxed areas are further magnified. Quantification of IHC results in tumors is shown to the right. **B**, Representative IHC of indicated proteins in liver from (**A**). Boxed areas are further magnified. **C**, Quantification of Ki67 staining in (**B**). **D**, Percentage of mice with PDAC liver metastases in (**A**). **E**, IB analysis of the indicated proteins in PANC-1 and KC6141 cells -/+ IL-8/CXCL2 or IL-17 for 5 days. **F**, Representative DNMT3B staining in pancreata from (**A**). Boxed areas show higher magnification. Image J quantitation of staining intensity is shown at the bottom. Data in (**A**, **F**) (*n*=6 mice), and (**C**) (*n*=8 fields) are mean ± s.e.m. Statistical significance was assessed by one-way ANOVA with Tukey post hoc tests. Exact *P* values are shown in the Source Data. \**P* < 0.05, \*\**P* < 0.01, \*\*\*\**P* < 0.0001. Scale bars 100 μm.

## Methods

### Cell Culture

All cells were incubated at 37℃ in a humidified chamber with 5% CO_2_. MIA PaCa-2, PANC-1, HPAC, SW1990, HEK293T, UN-KPC-960 (KPC), UN-KC-6141 (KC) cells, WT and R/R fibroblasts were maintained in DMEM (Invitrogen) supplemented with 10% fetal bovine serum (FBS) (Gibco). 1305 primary human PDAC cells, BxPC3 and AsPC1 cells were maintained in RPMI (Gibco) supplemented with 20% (1305) or 10% (BxPC3 and AsPC1) FBS and 1 mM sodium pyruvate (Corning). MIA PaCa-2 cells were purchased from ATCC. HEK293T and BxPC3 cells were provided by Dr. Michael Karin (SBP Medical Discovery Institute). HPAC, PANC-1 and SW1990 cells were purchased from Shanghai Yuchun Biology (Shanghai, China). KPC and KC cells were provided by Dr. Surinder K. Batra (University of Nebraska Medical Center). WT and R/R fibroblasts were generated as before (5). 1305 primary human PDAC cells were generated by Dr. Andrew M. Lowy (University of California San Diego) from a human PDAC PDX. All media were supplemented with penicillin (100 U/ml) and streptomycin (100 μg/ml). All cells were partially authenticated by visual morphology. WT and R/R fibroblasts were partially authenticated by ECM production and Col-I cleavage. All cell lines were routinely tested for mycoplasma contamination. Low glucose (LG) medium: glucose-free DMEM medium was supplemented with 0.5 mM glucose, 10% dialyzed FBS and 25 mM HEPES.

### Plasmids and transduction

For gene ablations, the target cDNA sequences of mouse (m) *Ddr1* (5’-GTAACGCAA CCGATAGCTTC-3’), and *Fbxw2* (5’-CCCCACTCAGACTAATCAGG-3’), and human (h) *FBXW2* (5’-CGTCTCTAAACAGTGGAATA-3’) were cloned into lentiCRISPR v2 -Blast vector or lentiCRISPR v2-puro vector, respectively, using BsmBI. For gene knockdowns (KD), the target cDNA sequences of h*DDR1* (5’-CCTATACGTTTCTGTGGAGTA-3’), h*FBXW2* (5’-GAACCAAAGTTCTCACCTAAT-3’), h*DNMT1* (5’-GAG GTTCGCTTATCAACTAAT-3’), h*DNMT3B* (5’-AGCCTAACACGGTGCTCATTT-3’), m*Ddr1* (5’-CGTCTCCATACTTGCCCATTC-3’), m*Fbxw2* (5’-GAACCAAAGTTCT CACCTAAA-3), m*Dnmt1* (5’-ACCAAGCTGTGTAGTACTTTG-3’), and m*Dnmt3b* (5’ -CCAGGAGTATTTGAAGATGAT-3’) were cloned into pLKO.1-puro vector. pLKO.1-puro-hTFAM (TRCN0000016095), pLKO.1-puro-hAMPKα1 (TRCN00000 00861), and pLKO.1-puro-mAmpkα1 (TRCN0000024000) were obtained from Sigma. pCDH-CMV-MCS-EF1-puro-Col1a1-6XHis, pCDH-CMV-MCS-EF1-puro-3/4Col1a1 -6XHis, pCDH-CMV-MCS-EF1-puro-1/4Col1a1-6XHis, pCDH-CMV-MCS-EF1-puro-Col1a1-R/R-6XHis were made by Sangon Biotech (Shanghai, China). pLVX-IRES-puro-Col1a2-HA, pLVX-IRES-puro-3/4Col1a2-HA, pLVX-IRES-puro-1/4 Col1a2-HA, pLVX-IRES-puro-Col1a2-R/R-HA, pLVX-IRES-puro-DDR1-WT-HA, pLVX-IRES-puro-DDR1-T519A-HA, pLVX-IRES -puro-T519E-HA, pLVX-IRES-puro-DDR1-K663R-HA, and pcDNA3.1-puro-FBXW2-HA, pcDNA3.1-puro-FBXW2^ΔF^-HA, and pLVX-IRES-puro-His-HA-ub, pLVX-IRES-puro-DNMT1-HA, pLVX-IRES-puro-DNMT3B-HA were made using ClonExpress Ultra One Step Cloning Kit V2-C116 (Vazyme, China). Flag-tagged Cullin 1, Cullin 2, Cullin 3, Cullin 4A, Cullin 4B, Cullin 5, and Cullin 7, β-TrCP1, SKP2, FBXW2, FBXW4, FBXW5, FBXW7, FBXW8, FBXO2, FBXO4, FBXL3, and FBXL5 cDNAs were cloned into pIRES2-puro vector.

For the system in which TET2 specifically demethylates the *FBXW2* DNA promoter, dCas9-TET2 catalytic domain (CD) fusion proteins were generated by cloning the TET2 CD cDNA into the C-terminus of the catalytically inactive Cas9 (dCas9) in the pLV-U6-MCS-sgRNA-dCas9-P2A-PuroR vector using the ClonExpress Ultra One Step Cloning Kit V2. The sgRNAs targeting the human *FBXW2* promoter (g1: GAGGAAACAGGGCATGAGAC; g2: CAGGTGAAGTGATTTGCCCA; g3: CAAG GTCACCCAGCAACAAG) or the sgRNAs targeting the mouse *Fbxw2* promoter (g1: CTGATATGTAACAAGCGCTT; g2: GTTCGGCCCTTTTCCACAGT; g3: GAGCC GCTTCCGCCCGACGC) were designed using DBCLS (https://crispr.dbcls.jp/) and cloned into the pLV-U6-MCS-sgRNA-dCas9-TET2 CD-P2A-PuroR vector (dCas9-TET2^sgFBXW2^ or dCas9-TET2^sgFbxw2^) with the ClonExpress Ultra One Step Cloning Kit V2.

Lentiviral particles were generated as before (8). PANC-1, HPAC, SW1990, 1305, KPC, or KC cells were transduced by combining 1 ml of viral particle-containing medium with 8 μg/ml polybrene. The cells were fed 8 h later with fresh medium and selection was initiated 48 h after transduction using 1.25 μg/ml puromycin or 10 μg/ml blasticidin.

### Mice

*Pdx1-cre*, *LSL-Kras^G12D+/-^*, *LSL-Tp53^R172H+/-^*, *Fbxw2^f/f^*, *Ddr1^f/f^*, and C57BL/6 mice were obtained at 6 weeks of age from Jiangsu Jicui Yaokang Biotechnology Co., Ltd. *Col1a1^+/+^*(Col-I^WT^) or *Col1a1^r/r^* (Col-I^r/r^) mice on a C57BL/6 background were obtained from Dr. David Brenner at UCSD and were previously described(5). *Pdx1- Cre*, *LSL-Kras^G12D^*, *Tp53^R172H+/-^*, *Fbxw2^f/f^*, and *Ddr1^f/f^* mice were interbred as needed to obtain the compound mutants *Pdx1-Cre*;*LSL-Kras*^G12D^ (termed *Kras^G12D^*), *Pdx1-Cre*;*LSL-Kras*^G12D^;*Fbxw2^f/f^* (termed *Kras^G12D^;Fbxw2^ΔPEC^*), *Pdx1-Cre*;*LSL-Kras*^G12D^; *Fbxw2^f/f^*;*Ddr1^f/f^*(termed *Kras^G12D^;Fbxw2^ΔPEC^;Ddr1^ΔPEC^*), *Pdx1-Cre*;*LSL-Kras*^G12D^; *LSL*-*Tp53^R172H+/-^* (termed *Kras^G12D^;Tp53^R172H^*), *Pdx1-Cre*;*LSL-Kras*^G12D^;*LSL*- *Tp53^R172H+/-^*;*Fbxw2^f/f^*(termed *Kras^G12D^;Tp53^R172H^;Fbxw2^ΔPEC^*), and *Pdx1-Cre*;*LSL-Kras*^G12D^;*LSL*-*Tp53^R172H+/-^*;*Fbxw2^f/f^*;*Ddr1^f/f^*(termed *Kras^G12D^;Tp53^R172H^;Fbxw2^ΔPEC^; Ddr1^ΔPEC^*). Mice matched for age and gender were randomly allocated to different experimental groups based on their genotypes. No sample size pre-estimation was performed but as many mice per group as possible were used to minimize type Ι/II errors. Both males and females were used unless otherwise stated. Blinding of mice was not performed except for IHC analysis. All mice were maintained in filter-topped cages on autoclaved food and water in constant temperature, humidity and pathogen-free controlled environment (23 ± 2°C, 50-60%) with a standard 12 h light/12 h dark cycle. Experiments were performed in accordance with Fudan University Institutional Animal Care and Use Committee guidelines and regulations. Dr. Su’s Animal Protocol DSF-2023-003 was approved by the Fudan University Institutional Animal Care and Use Committee. The number of mice per experiment and their age are indicated in the figure legends.

### Orthotopic and intrasplenic PDAC cell implantation

Parental, FBXW2^KD^, DDR1^KD^, DDR1/FBXW2^DKD^, DDR1^T519A^, or DDR1^K663R^ KPC or KC cells -/+ CXCL2 pretreatment or dCas9-TET2^sgFbxw2^ transfection were orthotopically injected into 2-month-old C57BL/6, Col-I^WT^ or Col-I^r/r^ mice as described(5). Following surgery, mice were given buprenorphine subcutaneously at a dose of 0.05-0.1 mg/kg every 4-6 h for 12 h and then every 6-8 h for 3 additional days. 5-AzC treatment (2.5 mg/kg daily i.p. injection) was initiated on day 3 post-transplantation. Mice were analyzed 3 weeks after transplantation.

### Human specimens

93 human PDAC specimens were acquired from patients diagnosed with PDAC between Jan. 2016 and Dec. 2022 at The First Affiliated Hospital of Anhui Medical University (Hefei, Anhui, China). All patients received standard surgical resection without chemotherapy before surgery. Paraffin embedded tissues were processed by a pathologist after surgical resection and confirmed as PDAC prior to further investigation. Overall survival duration was defined as the time from date of diagnosis to that of death or last known follow-up examination. The study was approved by the Institutional Ethics Committee of The First Affiliated Hospital of Anhui Medical University with IRB # PJ 2025-11-74. Informed consent for tissue analysis was obtained before surgery. All research was performed in compliance with government policies and the Helsinki declaration.

### Human PDAC organoids

Patient-derived fresh tissue samples were collected with written informed patient consent with the approval of the Institutional Ethics Committee (IRB # PJ 2025-11-74) at The First Affiliated Hospital of Anhui Medical University. Wash fresh tissue ≥3 times with a minimum of 10 volumes of wash solution (Advanced DMEM supplemented with 2% FBS, 2 mM EDTA and 10.5 μM Y-27632) to remove residual blood and interstitial fluid. Transfer the tissue to a 6-cm dish, add 1-2 ml of tissue dissociation solution and mince into fragments (<1 mm) using sterile scissors and surgical blades, removing any visible adipose tissue. Transfer the minced tissue to a sterile 100-ml Erlenmeyer flask containing ≥15 volumes of tissue dissociation solution [Advanced DMEM supplemented with 5 mg/ml collagenase II (Gibco), 1 mg/ml dispase (Yeason) and 0.1 mg/ml DNase I (Gibco)], and incubate at 37 °C with agitation at 120 rpm for 1 h. Filter the suspension through a 70-μm cell strainer, centrifuge the filtrate at 300g for 5 min and discard the supernatant. Wash the pellet by resuspending in wash solution followed by centrifugation (300g, 5 min); repeat ≥3 times. Resuspend the final pellet in 3 ml wash solution and filter through a 40-μm strainer. Centrifuge the filtrate (300g, 5 min) and resuspend the pellet in ≥5 volumes of Matrigel. Plate 50-μl droplets into 24- or 48-well plates, solidify at 37 °C for 30 min and overlay each droplet with 500 μl NGC Organoid Human Cancer Organoid Culture Medium: Pancreatic Cancer Organoid (D1Med). Refresh the medium every 2-3 days.

When organoids reach 300-500 μm in diameter, passage by adding ≥10 volumes of ice-cold PBS (with DNase) per droplet to dissolve Matrigel. Pipette gently, incubate on ice for 15 min, centrifuge (300g, 5 min), discard the supernatant and repeat dissociation-centrifugation three times to remove residual Matrigel. Digest organoids with TrypLE in a 37 °C water bath for 10-15 min, neutralize with an equal volume of wash solution and centrifuge (300g, 5 min). Repeat centrifugation three times to obtain single-cell suspensions for replating or cryopreservation. Replate single cells or clusters using the initial Matrigel-embedding procedure. For cryopreservation, suspend cells in Organoid Cryopreservation Solution (D1Med) and store in liquid nitrogen; repeat the dissociation-digestion sequence after thawing.

### Immunohistochemistry

Pancreata or livers were surgically removed, fixed in 4% paraformaldehyde in PBS and embedded in paraffin. 5 μm sections were prepared and stained with hematoxylin and eosin (H&E) or Masson’s trichrome (Masson). IHC was performed as before (8). Slides were scanned using a Motic EasyScanner with MoticEasyScanner software (MOTIC, China).

IHC scoring was performed as before (8). Negative and weak were viewed as low expression level and intermediate and strong were viewed as high expression level. For cases with tumors with two satisfactory cores, the results were averaged; for cases with tumors with one poor-quality core, results were based on the interpretable core. Based on this evaluation system, Chi-square test was used to estimate the association between FBXW2-Col-I-DDR1/pDDR1^T519^-NRF2 signaling proteins staining intensities. The number of evaluated cases for each different staining in PDAC tissues and the scoring summary are indicated (Figure S4b).

### Extracellular matrix preparation

WT or R/R fibroblasts were seeded on 6/12/96-well plates. One day after plating, cells were switched into DMEM (with pyruvate) with 10% dialyzed FBS supplemented with 100 μM vitamin C. Cells were cultured for 5 days with media renewal every 24 h. Then fibroblasts were removed by washing in 1 ml or 500 μl or 100 μl per well PBS with 0.5% (v/v) Triton X-100 and 20 mM NH_4_OH. The ECM was washed five times with PBS before cancer cell plating. The following day, cancer cells were switched into the indicated medium for 24 or 72 h.

### Cell imaging

Cells were cultured on coverslips coated -/+ ECM and fixed in 4% PFA for 10 min at room temperature. Macropinosome visualization in cells and tissues, and immunostaining, were performed as previously (8). Images were captured and analyzed using a Leica SP8 STED 3X confocal microscope with Leica Application Suite AF 3.8.1.26810 software (Leica, Germany).

### Immunoblot and immunoprecipitation

Preparation of protein samples from cells and tissues, immunoblot and immunoprecipitation were performed as before (8, 38). Immunoreactive bands were detected by Tanon 5200S, automatic X-ray film processor, or a KwikQuant Imager.

### ELISA assay for collagen-DDR1 interaction

ELISA assay for collagen-DDR1 interaction was described as before (39). Recombinant intact human collagen type I (iCol-I, R/R Col-I) or 3/4 Col-I was diluted to 20 ng/μl in 0.05 M carbonate-bicarbonate coating buffer (pH 9.6) and 50 µL/well was added to a high-binding 96-well plate (Corning). Plates were incubated overnight at 4 °C, washed three times with PBST and blocked with 50 µL blocking buffer (1% BSA-PBST) for 1 h at room temperature. After washing, 50 µL serial dilutions of purified flag-tagged human DDR1 (0-30 nM) was added to the collagen-coated wells. Plates were incubated for 1.5 h at 25 °C with gentle shaking, then washed five times with PBST. Bound DDR1 was detected by incubation with anti-Flag monoclonal antibody overnight at 4 °C followed by HRP-conjugated secondary antibody for 1h at room temperature. Following a final series of five washes with PBST, 50 µL of TMB substrate (Beyotime, P0208) was added and color developed for 2-5 min in the dark; the reaction was stopped with 50 µL 1 M H_2_SO_4_ (Sangon Biotech, K124FA000) and absorbance was measured at 450 nm using a microplate reader with Skanlt RE 7.0.1 software (Thermo Scientific™ Varioskan™ LUX, USA).

### Real-time PCR analysis

Total RNA and DNA were extracted using Trizol (RK30129, ABclonal). RNA was reverse transcribed using a ABScript GII Reverse Transcriptase (ABclonal). Real-time PCR was performed as described (8). Primers were obtained from the NIH Primer-BLAST (https://www.ncbi.nlm.nih.gov/tools/primer-blast/index.cgi?LINK_LOC= BlastHome) were as follows: h*DDR1* F: 5’-GAAGTTCACGACTGCGAGTG-3’; h*DDR1* R: 5’-TCGGACCCATCTTCCCTCTC-3’; h*FBXW2* F: 5’-GGCGCTCGCTCA GGTAAAT-3’; h*FBXW2* R: 5’-TCTGCAAGTCCGTCAGAGAA-3’; m*Fbxw2* F: 5’-GC TTCGGTTTTTCCTAGCCG-3’; m*Fbxw2* R: 5’-GGTCCAGAGTTTCATTTTTCTGT T-3’; h*ACTB* F: 5’-TCACCCACACTGTGCCCATCTAC-3’; h*ACTB* R: 5’-GGAACC GCTCATTGCCAATG-3’; m*Actb* F: 5’-TTCCTTCTTGGGTATGGAAT-3’; m*Actb* R: 5’-GAGCAATGATCTTGATCTTC-3’.

### Cell viability assay

Parental, FBXW2^KD^, DNMT1^KD^, DNMT3B^KD^, DNMT1/DNMT3B^DKD^, DNMT3B /FBXW2^DKD^, DDR1/FBXW2^DKD^, DDR1^WT^, DDR1^T519A^, DDR1^T519E^, or DDR1^K663R^ cells were plated in 96-well plates coated -/+ ECM at a density of 3000 cells (PANC-1, HPAC), 1500 cells (KPC or KC), 10000 cells (HPAC or SW1990) pretreated with or without IL8 (human cell) or CXCL2 (mouse cell) per well and incubated overnight prior to treatment. Then cells were cultured in the presence of complete medium, or LG with or without treatment of IL8 or CXCL2, or 5-AzC, or dCas9-TET2^sgFBXW2^ transfection for the period indicated in the figure legends. Cell viability was determined with Cell Counting Kit-8 assay (Glpbio). Optical density was read at 450 nm and analyzed using a microplate reader with Skanlt RE 7.0.1 software. For all experiments, culture medium was replaced every 24 h.

### ATP measurements

Cells were grown on 96-well plates coated -/+ indicated ECM for 24 or 72 h. Then cell number was measured. Intracellular ATP was determined with Varioskan^TM^ LUX (Thermo Scientific^TM^, USA) according to manufacturer’s protocol. Finally, luminescence was measured and normalized to cell number.

### Protein-Protein Docking

The amino acid sequences of human Col-I (composed of two COL1A1 chains and one COL1A2 chain) and DDR1 were retrieved from the UniProt database (UniProtKB; www.uniprot.org). Structural models of these proteins were predicted using AlphaFold3 (40) (https://alphafoldserver.com/). The predicted structures of iCol-I, cCol-I, and DDR1 were then subjected to protein-protein docking using the HDOCK server (http://hdock.phys.hust.edu.cn/), an integrative docking platform that combines template-based modeling with ab initio free docking (41). The top-ranked docking models were selected according to the HDOCK scoring function. The resulting complex structures were analyzed and visualized using PyMOL (version 3.1.6.1).

### HPLC-MS/MS

To identify phosphorylation or ubiquitination sites on DDR1 by mass spectrometry, gel bands corresponding to phosphorylated DDR1-Flag or endogenous DDR1 were excised and subjected to in-gel digestion. Peptides were separated on a nanoElute liquid chromatography system (Bruker Daltonics) equipped with a 250 mm × 75 μm analytical column. The mobile phases were 0.1% formic acid in water (A) and 0.1% formic acid in acetonitrile (B). Peptide elution was performed at a flow rate of 300 nL/min using a 1-hour gradient: solvent B was increased from 2% to 22% over 45 minutes, to 37% over 5 minutes, and then to 80% over another 5 minutes, followed by a 5-minute wash at 80% B.

Mass spectrometry analysis was carried out on a TIMS-TOF Pro instrument (Bruker Daltonics) equipped with a nano-electrospray ion source. The ion scan range was set to 100-1700 m/z, and the TIMS range was 0.7-1.3 Vs/cm^2^ with a 1.16-s cycle time (one MS scan followed by ten PASEF MS/MS scans). The intensity threshold was 5000, with ion accumulation and release times of 100 ms. The source voltage was 1500 V, with an auxiliary gas flow of 3 L/min and a source temperature of 180°C.

Raw MS data were processed using PEAKS Online (Bioinformatics Solutions Inc., version 1.7) against the human SwissProt database (20,331 entries). Search parameters included a precursor mass tolerance of 20 ppm and a fragment mass tolerance of 0.05 Da, semi-tryptic specificity, carbamidomethylation of cysteine as a fixed modification, and variable modifications including N-terminal acetylation, methionine oxidation, and asparagine/glutamine deamidation. Protein peak areas were extracted for subsequent statistical analysis.

### Single-cell ATAC sequencing (scATAC-seq) analysis

scATAC-seq data from PDAC specimens were obtained from the Human Tumor Atlas Network (HTAN) data portal (https://data.humantumoratlas.org/). The data were analyzed in R (v4.4.1) using the Seurat (42) (v5.1.0) and Signac (43) (v1.14) packages. Quality control (QC) was performed to retain high-quality nuclei according to the following criteria: 2,000 < nCount_ATAC < 30,000; blacklisted region ratio < 0.02; nucleosome signal < 4; transcription start site (TSS) enrichment > 2; and fraction of reads in peaks (pct_reads_in_peaks) > 15%. Only cells meeting all QC thresholds were included in downstream analyses.

Cell type annotation was achieved by integrating the scATAC-seq data with matched scRNA-seq data from the same cohort using Seurat’s label transfer method. UMAP was applied for two-dimensional visualization of the integrated dataset. To explore locus-specific chromatin accessibility, the CoveragePlot function in Signac was used to visualize ATAC-seq signal profiles in normal epithelial and malignant epithelial cells at the genomic region encompassing the FBXW2 locus. Differentially accessible regions (DARs) were identified and highlighted accordingly (chr9: 120,792,300-120,792,700, hg38/GRCh38).

### Bisulfite sequencing analysis

Genomic DNA were extracted from the cells indicated in the figure legends using a genomic DNA extraction kit (QIAGEN). 2 μg DNA was used to convert unmethylated cytosines to uracil according to the manufacturer’s protocol (Beyotime, D0068S). Then the *FBXW2* promoter region was amplified by a BisPCR^2^ protocol described as before (44). The *FBXW2* promoter region was amplified by the following primers: h*FBXW2*: F: 5’-ACACTCTTTCCCTACACGACGCTCTTCCGATCTTTGTTTTTAGGTTTTT TGTTTT-3’; R’5’-GTGACTGGAGTTCAGACGTGTGCTCTTCCGATCTACTATTTTTCTTTCCCTACAAAC-3’; m*Fbxw2* F: 5’-ACACTCTTTCCCTACACGACGCTCTTCCGATCTTTTGTTTAGGATTATAAGGTGTAAG-3’; R: 5’-GTGACTGGAG TTCAGACGTGTGCTCTTCCGATCTTAAAAATTCCTATCCAACCC-3’. Then the PCR products were used to do the second PCR to prepare DNA sequencing libraries by adding the different barcodes for distinct treated samples. All the PCR^2^ primers are same except the barcodes highlight by the underline in the reversed primers. The Primers for PCR as follows: F: 5’-AATGATACGGCGACCACCGAGATCTACACTCTTTCCCTACACGAC-3’; R: 5’-CAAGCAGAAGACGGCATACGAGAT<u>CGTG</u> <u>AT</u>GTGACTGGAGTTCAGACGTGT-3’; 5’-CAAGCAGAAGACGGCATACGAGA T<u>ACATCG</u>GTGACTGGAGTTCAGACGTGT-3’; 5’-CAAGCAGAAGACGGCATA CGAGAT<u>GCCTAA</u>GTGACTGGAGTTCAGACGTGT-3’; 5’-CAAGCAGAA GACG GCATACGAGAT<u>TGGTCA</u>GTGACTGGAGTTCAGACGTGT-3’; 5’-CAAGCAGA AGACGGCATACGAGAT<u>CACTGT</u>GTGACTGGAGTTCAGACGTGT-3’; 5’-CAA GCAGAAGACGGCATACGAGAT<u>ATTGGC</u>GTGACTGGAGTTCAGACGTGT-3’.

The PCR^2^ products were sent for DNA sequencing using Nanopore sequencing. Raw sequencing reads were initially processed to isolate target PCR amplicons based on unique primer sequences. Specifically, reads containing both the forward and reverse primer sequences were extracted using Seqkit (v2.10.1). The extracted reads were then assigned to individual samples according to their unique index barcodes ligated during library preparation. Sample-specific FASTQ files were subjected to adapter and quality trimming using Trim Galore (v0.6.10), configured for bisulfite-converted DNA. The cleaned reads were subsequently converted from FASTQ to FASTA format. Finally, the resulting FASTA files were submitted to QUMA (45) (Quantification Tool for Methylation Analysis; http://quma.cdb.riken.jp/) for bisulfite sequence alignment and methylation analysis.

### MeDIP-qPCR assay

Methylated DNA IP (MeDIP) was performed using the EpiQuik Methylated DNA Immunoprecipitation Kit (Epigentek, P-2019-48). In brief, DNA was extracted from SW1990, HPAC, BxPC3 cells and human PDAC organoids, sheared and added into microwells coated with IgG and 5-methylcytosine antibodies. DNA was released from the antibody-DNA complex and purified through Fast-Spin Columns. qPCR assays were then performed with eluted DNA with the primers as following: h*FBXW2*’F: 5’-CCTTGCTCTCGGAATCGTGT-3’; R: 5’-GCGTGCCTAGTAGCTGTGAA-3’; m*FBXW2*’F: 5’-CGCGCAGTTAGGGATTTGTTC-3’; R: 5’-GGGCTAAGGCTC CAAATCCA-3’.

### Statistics and reproducibility

Macropinosomes or mitochondria were quantified using the ‘Analyze Particles’ feature in Image J (National Institutes of Health). Macropinocytotic uptake index (46) or mitochondria number were computed by macropinosome or mitochondria area in relation to total cell area for each field and then by determining the average across 6 fields. Tumor area (%) was quantified using the ‘Polygon’ and ‘Measure’ feature in Fiji Image J and was computed by tumor area in relation to total area for each field and then by determining the average across the indicated mice. Positive area of protein expression in tissues (%) was quantified using ‘Colour Deconvolution’, ‘H DAB’, and ‘Analyze Particles’ feature in Fiji Image J and was computed by protein positive area in relation to the tissue or tumor area for each field and then by determining the average across the indicated fields or mice. These measurements were done on randomly-selected fields of view. Two-tailed unpaired Student’s t test, one-way ANOVA with Tukey post hoc tests, or Mann-Whitney test, Kruskal-Wallis tests with Dunn post hoc test based on data normality distribution was performed for statistical analysis using GraphPad Prism software. Data are presented as mean ± SEM. Kaplan-Meier survival curves were analyzed by log rank test. Binding data were plotted as mean ± s.e.m. and analyzed in GraphPad Prism using nonlinear regression (one-site total binding) to derive apparent binding parameters and Kd estimates. Statistical comparisons were performed as indicated in the figure legends.

Statistical correlation between FBXW2-Col-I-DDR1-NRF2 signaling proteins in human PDAC specimens was determined by two-tailed Chi-square test. (\*\*\*\**P* < 0.0001, \*\*\**P* < 0.001, \*\**P* < 0.01, \**P* < 0.05). All experiments except IHC analysis of 93 human specimens were repeated at least three times.

## Data availability

Graph data are provided in Source data. All raw image data including immunostaining, immunoblot, IHC, H&E and Masson staining were uploaded to Mendeley Data (10.17632/v3wzrnnvkr.1).

## Acknowledgements

We thank members of the H.S. and M.K. laboratories for discussions; the Core Facility of Shanghai Medical College, Fudan University for mass spectrometry and microscopy services. Research was supported by grants from the National Natural Science Foundation of China (Grant Nos. 82372884, 82573758, 2022hwyq29 to H.S., 82372644 to F.Y., 82120108012 and 81930086 to B.S.); Noncommunicable Chronic Diseases-National Science and Technology Major Project (2026ZD0555800 to H.S.); the Shanghai Natural Science Foundation (Grant No. 23ZR1413600 to H.S.); the Fund of Fudan University and Cao’ejiang Basic Research (Grant No. 24FCA02 to H.S.); the Fund of Anhui Province Higher Education Outstanding Young Researcher (Grant No. 2024AH020007 to F.Y.); Padres Pedal the Cause/C3 (PPTC2018 to M.K.); the USA NIH (R01CA211794, R37AI043477, P01DK098108, U01CA274295 and U01AA027681 to M.K.); Clinical Research Special Project of Anhui Provincial Department of Science and Technology (202204295107020008 to B.S.); and Research Program of Anhui Provincial Department of Education (B.S.); Additional support was provided by Ride the Point (M.K.). Author contributions H.S., M.K. and F.Y. conceived the project. H.S. designed the study and F.Y. and H.S. performed most experiments. F.Y., H.S., B.L., X.P. and R.L. performed IHC analysis of human and mouse PDAC. F.Y., H.S., B.L., K.W., and C.W. performed immunoblotting and qPCR analysis. H.S., F.Y., and B.L. performed orthotopic PDAC cell implantations. Z.Y. prepared human PDAC organoids. Y.Y. performed bioinformatics analysis. B.S. collected human PDAC tissue and supervised and supported F.Y. and Z.Y.. H.S., M.K., and F.Y. wrote the manuscript, with all authors contributing and providing feedback and advice.

